# Gene expression based inference of drug resistance in cancer

**DOI:** 10.1101/2021.11.17.468905

**Authors:** Smriti Chawla, Anja Rockstroh, Melanie Lehman, Ellca Rather, Atishay Jain, Anuneet Anand, Apoorva Gupta, Namrata Bhattacharya, Sarita Poonia, Priyadarshini Rai, Nirjhar Das, Angshul Majumdar, Jayadeva, Gaurav Ahuja, Brett G. Hollier, Colleen C. Nelson, Debarka Sengupta

## Abstract

Inter and intra-tumoral heterogeneity are major stumbling blocks in the treatment of cancer and are responsible for imparting differential drug responses in cancer patients. Recently, the availability of large-scale drug screening datasets has provided an opportunity for predicting appropriate patient-tailored therapies by employing machine learning approaches. In this study, we report a predictive modeling approach to infer treatment response in cancers using gene expression data. In particular, we demonstrate the benefits of considering integrated chemogenomics approach, utilizing the molecular drug descriptors and pathway activity information as opposed to gene expression levels. We performed extensive validation of our approach on tissue-derived single-cell and bulk expression data. Further, we constructed several prostate cancer cell lines and xenografts, exposed to differential treatment conditions to assess the predictability of the outcomes. Our approach was further assessed on pan-cancer RNA-sequencing data from The Cancer Genome Atlas (TCGA) archives, as well as an independent clinical trial study describing the treatment journey of three melanoma patients. To summarise, we benchmarked the proposed approach on cancer RNA-seq data, obtained from cell lines, xenografts, as well as humans. We concluded that pathway-activity patterns in cancer cells are reasonably indicative of drug resistance, and therefore can be leveraged in personalized treatment recommendations.

## Introduction

Cancer is a highly complex disease exhibiting varying degrees of genetic and phenotypic heterogeneity within individuals. Despite the apparent overall improvement in prognosis, response to existing therapies (chemotherapies and immunotherapies) are often unpredictable. This is primarily attributable to the clonal diversity of cancer cells and associated phenotypically altered non-malignant cells in the tumor microenvironment. These pose a substantial hindrance to the optimal therapeutic management of the disease^1,2^. As a rescue, presently, therapies that work by targeting specific molecules or mutations are highly advocated. EGFR expression & mutations, KRAS mutations, BCR-ABL fusions, and HER2 overexpression are examples of popular therapeutic targets in cancer^3^. Such therapies are frequently rendered futile or counterproductive due to the unavailability of known molecular targets in patient tumors. As such, it’s concluded that drug targets or mutational status alone are incomprehensive as indicators for targeted therapies^4,5^. Furthermore, administering a targeted therapy without considering drug resistance as a consequence may worsen patient survival. As such, early inference of drug response, based on pretreatment molecular portraits of cancer has become a necessity^6,2^.

In recent years, the availability of large-scale pharmacogenomic databases has propelled predictive personalized oncology research^2^. Cancer Cell Line Encyclopedia (CCLE)^7^ and Genomics of Drug Sensitivity in Cancer (GDSC)^8^ are noteworthy among these^9^. These studies embody an expansive knowledgebase comprising high-throughput screening studies entailing more than 1000 cell lines with hundreds of anticancer drugs^2^. Concurrently, The Cancer Genome Atlas (TCGA)^10^ serves as another rich database featuring gene expression profiles of primary tumors spanning multiple cancer types, with associated clinical meta-data and drug-response annotations. This wealth of data has enabled the modeling of drug response based on molecular profiles. Various machine learning methods have been proposed for drug response prediction in cancer. Jia et al. proposed a deep variational autoencoder for imputing drug response through compression of multiple genes into latent vectors in low dimensional space^9^. Ammad-Ud-Din, Muhammad, et al.^11^ reported a kernelized bayesian matrix factorization based drug response prediction through incorporating prior knowledge of pathway-drug associations. Another approach utilized matrix factorization with similarity regularization for drug response prediction in cell lines by employing chemical structures of drugs and gene expression profiles^12^. Chang and colleagues proposed CDRscan, a convolutional neural net that leverages mutational signatures for predicting drug effectiveness^13^. Sakellaropoulos, Theodore, et al. reported gene expression data based deep neural network for drug response prediction that outperforms ElasticNet and Random Forest^14^.

By carefully surveying these methods, we identified two critical scopes for improvement. First, the majority of the past studies do not consider structural properties of drugs as features (explanatory variables in predictive tasks). As a result, the machine learning models learn suboptimally and fail to make predictions on drugs that are not part of the training data. Secondly, gene expression levels are considered as independent variables, ignoring their pathway-specific combinatorial implication. Past studies have demonstrated the utility of using pathway enrichment scores for various downstream analyses, as opposed to gene expression values^15,16^. Notably, in our previous works, we have shown how pathway projection facilitates better modeling of biological processes^16,17^. As an added advantage, data integration based on pathway enrichment scores subsides batch-effects^18^. While single-cell RNA-seq (scRNA-seq) enables characterization of cellular heterogeneity in tumors, there’s very little visible effort to leverage this fine-grained molecular information to predict drug-response at sub-clonal resolution. This is primarily due to the fact that most available training data, as indicated above, are bulk expression profiles, and training on bulk RNA-seq and testing on scRNA-seq is expected to give rise to misleading predictions. Pathway projections of scRNA-seq/bulk RNA-seq profiles reasonably alleviate this problem. Notable in this regard is the work by Suphavilai, Chayaporn, et al.,^19^ that describes a drug response prediction approach in head and neck cancer, leveraging scRNA-seq profiles. The authors, however, did not explore the utility of drug descriptors to generalise the prediction model.

In this work, we used various machine learning and deep learning approaches (DNN) to train models for the prediction of drug-response in both *in vitro* and *in vivo* settings. We made use of publicly available high-throughput screening data (CCLE and GDSC) concerning a multitude of drug-cancer cell line pairs, as well as patient data from TCGA. We validated the cell line based model on multiple scRNA-seq datasets of established cancer cell lines.

Convinced by the reproducibility and overall performance of the cell-line model, we delved deeper into similar prediction tasks pertaining to in-house prostate cancer cell lines and animal models under differential treatment conditions. Prostate cancer is the most common malignancy in men, yet for patients presenting with advanced cancer, treatment options are limited. Androgen deprivation is an effective therapeutic strategy widely used in clinical practice. It exploits the unique dependence of prostate cancer on androgen signaling for growth and progression. Although androgen deprivation therapy is effective in most patients, the effect is only palliative and cancer cells eventually become resistant with the emergence of metastatic castration-resistant prostate cancer (CRPC). To date, only a limited number of agents have shown effectiveness and have been approved for the treatment of CRPC, yet their conferred survival benefit is only minimal, and acquired resistance to the drugs eventually emerges. Thus, appropriate drug selection and combination remain crucial in the dynamically cancer evolving landscape to derive maximum benefit for the patients^20,21,22^. We employed our internal bulk RNA-seq profiles of prostate cancer cell lines under different treatment conditions for further validation. To ensure the cross-sample utility of our model, we examined our LNCaP xenograft dataset mirroring *in vitro* treatments. For this, we utilized LNCaP xenografts from a prostate cancer progression study where the tumors were harvested at different stages, namely Pre-castration (PRE-CX), post castration (POST-CX), castration-resistant prostate cancer (CRPC), and while on enzalutamide (ENZ) treatment during the course of tumor progression. Our results revealed clinically and biologically relevant associations of drugs and pathways in terms of resistance and sensitivity. Along the same lines, we benchmarked the efficiency of the model, trained on patient samples from TCGA on RNA-seq profiles of pre-treatment, and matched post relapse drug-resistant BRAF mutant melanoma cancer patients. Our study connects a systematic drug response prediction pipeline with layered *in vitro* and *in vivo* validations involving cell lines, xenografts, and patients, which is the most important prerequisite for the clinical implementation of such approaches.

## Results

### Overview of approach

In this study, we model drug response using gene expression data in both *in vitro* and *in vivo* settings. For this, we leveraged Cancer Cell Line Encyclopedia (CCLE) cancer cell line gene expression RNA-seq bulk profiles. We formulated the prediction of drug response in terms of Z-score (LN IC50) as a regression task and constructed a suitable deep neural network (DNN) architecture. We performed rigorous assessment of drug response prediction in terms of LN IC50 and Z-score (LN IC50) on several datasets. Previous studies have reported use of IC50 for modelling drug response^23^. However, our analysis didn’t reveal biologically reasonable results with LN IC50 (data not shown). Hence, we resorted to Z-scores (LN IC50) for our analysis. First, we converted bulk RNA-seq gene expression profiles of cell lines from the CCLE project into pathway scores using GSVA^15^. Second, we integrated the cell line pathway scores matrix with the simplified molecular-input line-entry system (SMILES) embeddings (numerical descriptors of molecules) of the drugs. We sourced drug response information for the CCLE cell lines from the Genomics of Drug Sensitivity in Cancer (GDSC) database. The summary of the datasets used is provided in **Supplementary Fig.1** We found 550 cell lines common in both the databases screened against 173 unique molecular compounds for which SMILES notations were available. We retrieved SMILES of 173 compounds using PubChemPy^24^ and converted them into vector embeddings of size 100 using the SMILES2Vec python package^25^. The final matrix used for training the model constituted 80056 cell line-drug combinations in rows and 1429 features entailing 1329 pathway score vectors and vectors of size 100 each for molecular descriptors in columns.

These vectors form a set of explanatory variables to predict drug response as obtained from the screening experiment datasets (**Fig. 1a**). We used the open-source software Keras^26^ to build a DNN architecture (**Fig. 1b**). We evaluated the performance of the DNN model on the CCLE bulk RNA-seq cell lines test dataset and achieved a coefficient of determination (R^2^) = 0.60; Pearson’s correlation coefficient = 0.78; *P*-value p < 2.2e−16 (**Fig. 1c**). We then compared the performance of the DNN model with the other two commonly used machine learning models: random forest (RF) and elastic net that have been used by previous studies for drug response prediction^27,28,29^ **(Supplementary Fig. 2).** The DNN architecture outperformed the other two models. As a control, we also evaluated the performance of these three models using gene expression vectors for 500 genes (as opposed to pathway score matrix) selected based on the coefficient of variation square (CV^2^). We observed that in gene-based models DNN model is better in comparison to RF and elastic net **(Supplementary Fig. 3a, b)**. Further, our results revealed that the performance of pathway-based models is better than gene-based models in the case of the elastic net and DNN. These findings substantiate our choice of pathway scores over gene expression values. To ensure an unbiased evaluation of our DNN model, we applied our CCLE bulk RNA-seq model to external datasets including both bulk RNA-seq and scRNA-seq profiles. We used the Kinker, G. S. et al. scRNA-seq cell line dataset^30^. After processing, this dataset consisted of 17279 drug-cell line combinations entailing 116 cell lines and 173 drugs **(see methods)**. We were able to achieve a coefficient of determination (R^2^) = 0.42; Pearson’s correlation coefficient = 0.65; *P*-value p < 2.2e−16 (**Fig. 1d**). As expected, we did not observe a reasonable correlation in scRNA-seq profiles than we achieved on bulk RNA-seq profiles. Further, we benchmarked our model using another scRNA-seq dataset from a previously published study by Lee et al*.^31^* consisting of treatment-naive breast cancer cells (MDA-MB-231) and a population of cells that had gained sensitivity to paclitaxel. In this study, the metastatic MDA-MB-231 cells were exposed to a paclitaxel drug. Most of the cells died after 5 days of exposure. However, some of the residual cells, cultured in drug free medium after withdrawal of drug, recommenced proliferation and established clones. Interestingly, these cells became more sensitive to paclitaxel on reexposure. To verify that these cells are more sensitive to paclitaxel, we applied the DNN model to this dataset and found that our model predicted the paclitaxel-sensitive population to be more sensitive to paclitaxel in comparison to the treatment-naive population, corroborating with the original findings **(Fig.1e)**.

**Fig. 1.**
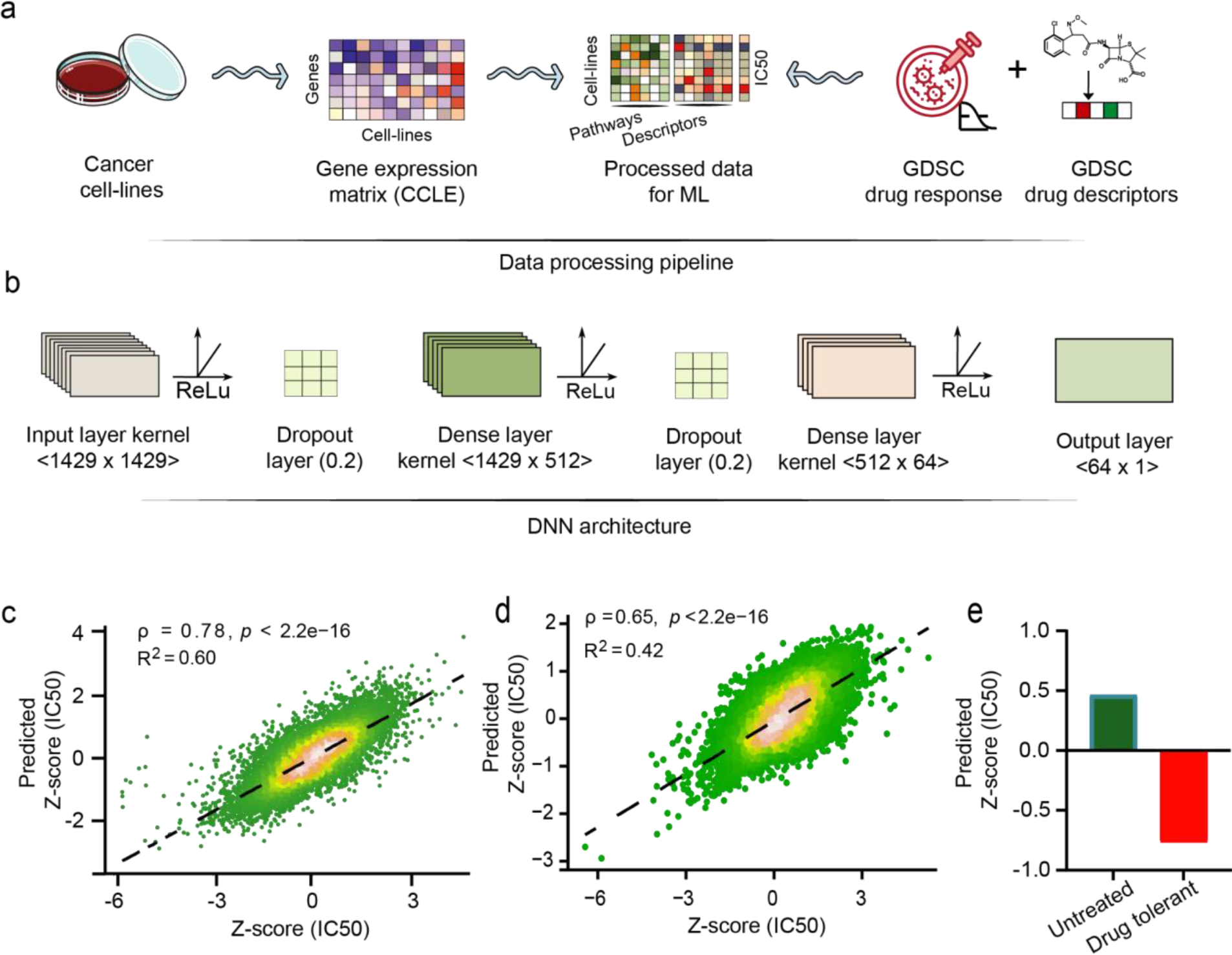
Illustration of the predictive analysis workflow. **a** Schematic workflow depicting data processing pipeline. The first step involves the processing of training data. The RNA-seq RSEM TPM genes expression profiles from Cancer Cell Line Encyclopedia (CCLE) were subjected to pathway score transformation using GSVA. This GSVA score matrix was integrated with the drug descriptors obtained in the form of SMILES embedding for each compound. **b** Model architecture. The second step was training of ML model on this data, comprising GSVA scores and drug descriptors as an explanatory variable set and Z-scores of the natural log-transformed IC50 values sourced from the GDSC database as the response variable. A deep neural network (DNN) from the Keras platform was used to perform the regression task of predicting drug response. **c** Scatter plot demonstrating model efficiency on the CCLE bulk RNA-seq test dataset in terms of observed vs predicted Z-score (LN IC50) measured by Pearson correlation (*ρ*) = 0.78 and coefficient of determination (R^2^) = 0.60. **d** Scatter plot demonstrating model efficiency on the Kinker, G. S. et al. scRNA-seq cell lines dataset comparing observed vs predicted Z-score (LN IC50) measured by Pearson correlation (*ρ*) = 0.65 and coefficient of determination (R^2^) = 0.42. **e** Evaluation of predicted drug response of paclitaxel in scRNA-seq MDA-MB-231 cells. Barplots depicting predicted response for paclitaxel in treatment-naive and paclitaxel sensitive cell population.

### Evaluation of models trained on CCLE bulk RNA-seq profiles using independent datasets

Despite therapeutics advances in prostate cancer, treatment options remain limited and emergence of treatment resistance poses significant challenges. Therefore, there is an unmet need for selection of optimal drug for prostate cancer treatment^32,33^. We independently validated our DNN model trained on CCLE bulk RNA-seq profiles using our in-house prostate cancer datasets. We applied the DNN model to bulk RNA-seq profiles of five untreated prostate cancer cell lines with two biological repeats of each cell line. We predicted drug responses for each of these ten samples for 155 drugs tested against prostate cancer cell lines in the GDSC database targeting various cellular pathways. Overall, we found the Androgen Receptor (AR) negative prostate cancer cell lines DU145 and PC3 were predicted to be more resistant to these therapeutic drugs, while the AR-positive cell lines LNCaP, DUCAP, and VCAP were more sensitive (**Fig. 2a**). Of these five cell lines, LNCaP cells were predicted to be most sensitive to these drugs (**Fig. 2b**). More specifically, the model predicted the sensitivity of LNCaP cells to PI3K/mTOR signaling pathway targeting drugs, in particular AKT inhibitors, afuresertib, and uprosertib **(Supplementary Fig. 4a)**. The LNCaP cells had high GSVA pathway scores for mTOR-related signaling, which may contribute to the predicted sensitivity to these AKT targeting drugs **(Supplementary Fig. 4b)**. When comparing the predicted Z-scores with the corresponding GDSC Z-scores, we observed a high Pearson correlation of 0.78 and 0.84 with a statistically significant *P*-value p < 2.2e−16 for the two biological replicates of LNCaP cells, respectively (**Fig. 2c**).

**Fig. 2.**
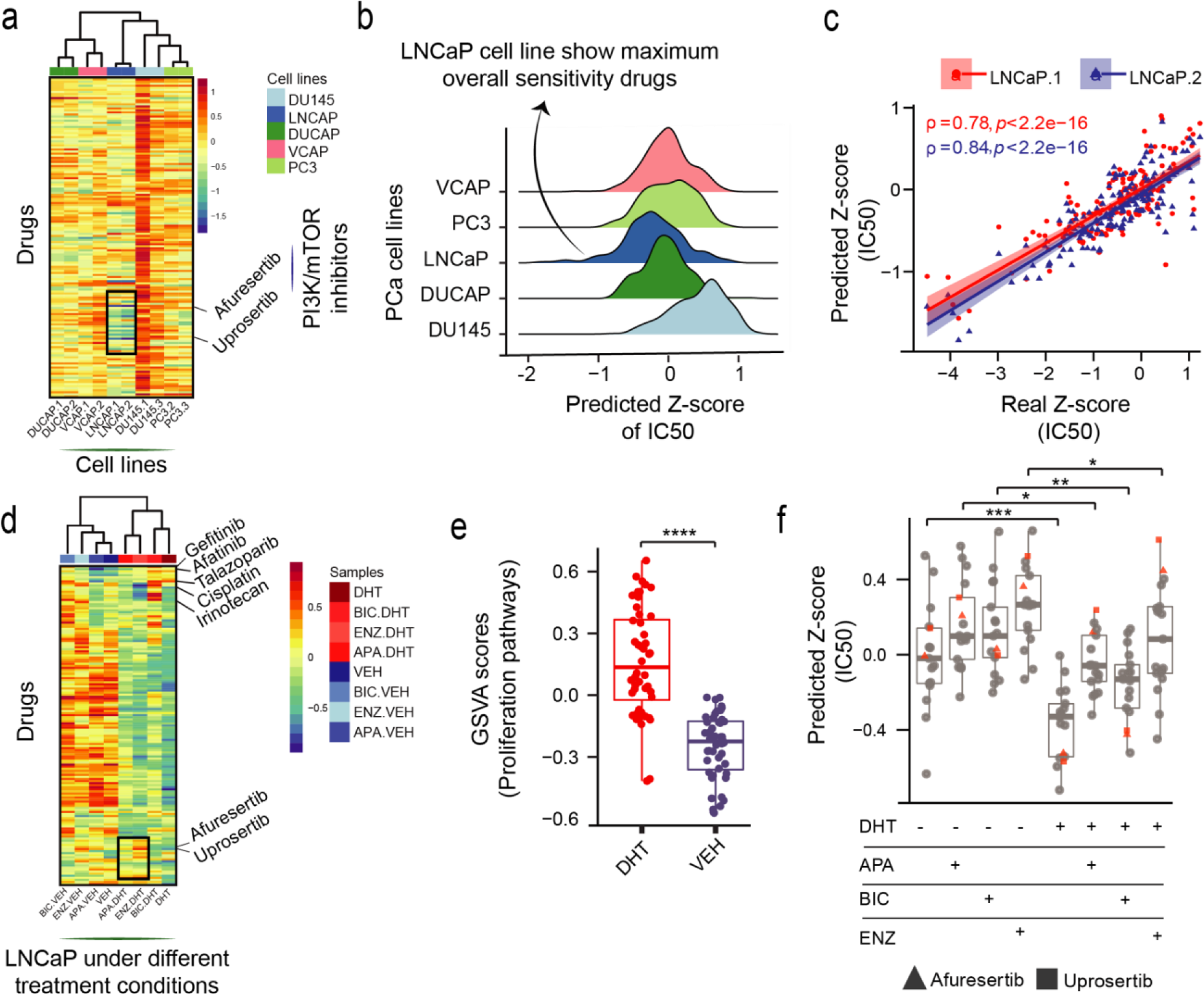
Analysis of drug response prediction in prostate cancer cell lines. **a** Heatmap showing predicted Z-scores (LN IC50) for 155 drugs across five prostate cancer baseline cell lines highlighting PI3K/mTOR signaling targeting drugs. The lower the Z-score, the more sensitive a sample is predicted to a drug. Color bars indicate cell lines type. **b** Ridgeplot showing overall distribution of predicted Z-scores (LN IC50) across five prostate cancer cell lines. **c** Scatterplot depicting Pearson correlation (*ρ*) between real and predicted Z-scores (IC50) for the LNCaP cell line. The line color indicated two biological replicates of the LNCaP cell line. **d** Heatmap of predicted Z-scores (LN IC50) of LNCaP cells in the presence and absence of androgens (DHT) and AR antagonists (ENZ, BIC, and APA) to 155 drugs. **e** Boxplots depicting the distribution of GSVA scores of proliferation-related pathways in the presence vs. absence of DHT (Asterisks denote Wilcoxon rank sum test *p-value*). **f** Boxplot of predicted Z-scores for drugs targeting PI3K/mTOR signaling across the treatment conditions (Asterisks denotes Wilcoxon rank sum test *p-value*).

We were further interested in testing how the drug response prediction was altered when LNCaP cells are cultured with androgen receptor (AR) agonist dihydrotestosterone (DHT) as compared to the vehicle control (VEH) in androgen deprived media conditions; and furthermore, how treatment with clinically approved AR antagonists bicalutamide (BIC), enzalutamide (ENZ), and apalutamide (APA) under these conditions affected the predicted sensitivity pattern. We used averaged GSVA scores of the biological replicates for the downstream analysis. Overall, cells cultured with DHT were predicted to be more sensitive to therapeutic drugs, while the cells cultured with no DHT and AR antagonists were more resistant **(Fig. 2d, Supplementary Fig. 5a)**. The cells cultured in the presence of DHT had high GSVA pathway scores of proliferation-associated pathways. Androgens are involved in stimulating prostate cancer cell proliferation^34,35^. This supports the notion that actively proliferating cells are more sensitive to particular anti-cancer drugs, while cells in a cytostatic or quiescent state are more resistant (**Fig. 2e**). Notably, the addition of AR antagonists in the presence of DHT did not fully reverse the predicted DHT conferred drug sensitivity. However, for a small subset of drugs mainly targeting the PI3K/mTOR signaling pathway, AR antagonists decreased the predicted drug sensitivity (**Fig 2f**). In particular, our DNN model predicted that treatment with ENZ in the presence of DHT compared to DHT alone rendered the cells more resistant to AKT inhibitors, in particular uprosertib and afuresertib (**Fig. 2d, f**). In the absence of androgens, ENZ and BIC treatment predicts increased resistance to drugs targeting EGFR signaling, with gefitinib and afatinib having the most profound effect. Whereas the addition of DHT alone increased drug resistance to EGFR targeting agents irrespective of the inclusion of AR antagonists **(Supplementary Fig. 5d)**.

Next, we evaluated our ability to predict drug response in LNCaP derived xenografts. Xenografts have emerged as useful *in vivo* cancer models for directly investigating therapeutic response and predicting anti-cancer drug response in patients with cancer of a similar phenotype. To this end, we used in-house data from LNCaP xenografts derived from a large and well-annotated prostate cancer progression study investigating responsiveness and subsequent resistance to therapies targeting the AR. LNCaP xenograft tumor establishment and initial growth are dependent on androgens in male mice (PRE-CX). Upon castration, AR activity and tumor growth is suppressed (POST-CX), however, this responsiveness to castration reproducibly gives way to castration-resistance (CRPC). Further treatment of CRPC with ENZ initially provides a therapeutic response (ENZ Sensitive; ENZS), however, resistance emerges in time (ENZ Resistant; ENZR) (**Fig. 3a**). Using our DNN model, we predicted the drug response for each of the 54 samples across this spectrum of successive therapeutic responsive and resistance states. The LNCaP xenograft tumor samples clustered into three main groups based on their overall predicted sensitivity to the 155 drugs in the analysis **(Fig. 3b, Supplementary fig. 6a)**. Cluster 1 samples had the most resistant tumors, which correlated to their lower proliferative index. Cluster 1 consisted of one CRPC sample and the remaining were ENZ-treated tumor samples (7 of the total 15 ENZR and 9 of 11 ENZS samples). In contrast, cluster 3 samples had the highest predicted overall sensitivity to the 155 drugs, which may be attributed to their higher proliferative index, indicated by higher GSVA pathway scores for cell proliferation-associated gene sets (**Fig. 3c, d**). Cluster 3 consisted of all PRE-CX mice tumors and subsets of the resistance types (5 of 10 POST-CX; 7 of 10 CRPC; and 4 of 15 ENZR samples). ENZR tumors were distributed across all three clusters, suggesting that resistance to ENZ occurs through different underlying mechanisms and more detailed analyses may uncover differential susceptibility to particular drugs in ENZR. This suggestion of multiple ENZ resistance mechanisms was strengthened by the multimodal distribution of predicted Z-scores for ENZR tumors compared to the uniform distribution for ENZS tumors (**Fig 3e**). In contrast to ENZS, ENZR samples were predicted to develop some level of sensitivity to a subset of drugs. ENZR samples tended to have higher GSVA scores of proliferation-associated pathways relative to ENZS samples however, this did not reach statistical significance (**Fig 3f**). ENZR tumors were predicted to be more sensitive to EGFR targeting drugs than any other tumor type in the study (**Fig 3g**), with sapitinib having the most profound effect **(Supplementary Fig S6b)**. While we achieved promising informative results on the drugs within our training set, we could also predict biologically relevant responses for drugs not included in the training set— APA, BIC, and ENZ. We observed sensitivity to AR antagonists in the PRE-CX, POST-CX and CRPC groups. Notably, we found responsive ENZS tumors from mice actively treated with ENZ were predicted to not benefit from additional AR antagonists. (**Fig 3h**).

**Fig. 3.**
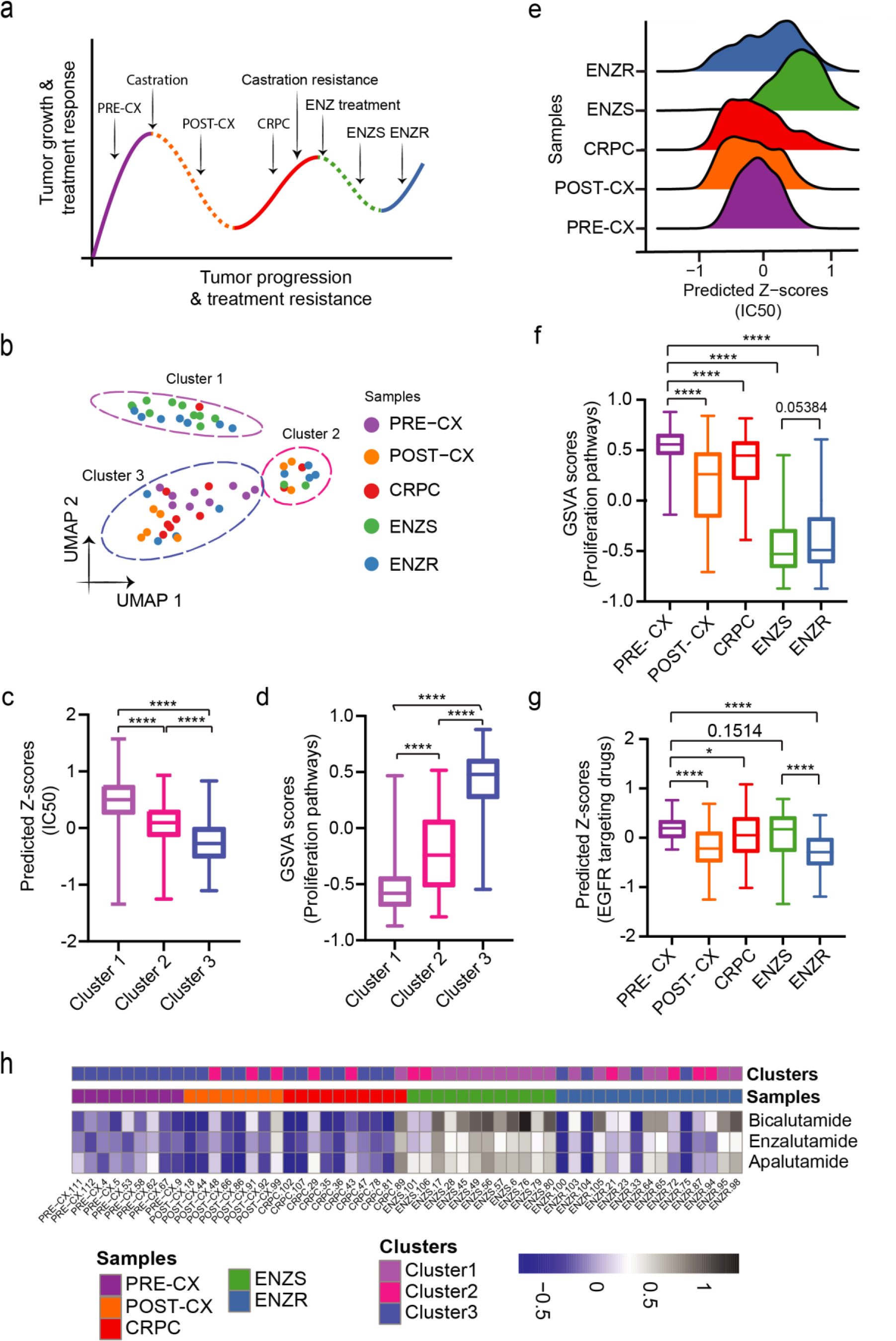
Analysis of drug response prediction in LNCaP derived xenografts. **a.** Overall schematics of the experimental setup of LNCaP xenograft-based prostate cancer progression study with indicated treatments, therapeutic response, and therapeutic resistance stages. Solid lines represent growth and treatment resistance; dotted lines are treatment responsiveness **b** UMAP based projections of predicted Z-scores showing three separate clusters. **c** Box plots depicting distribution predicted Z-score (LN IC50) across 3 clusters (Asterisks denote Wilcoxon rank sum test *p-value*). **d** Boxplots showing the distribution of GSVA scores of proliferation-related pathways across 3 clusters (Asterisks denote Wilcoxon rank sum test *p-value*). **e** Ridgeplot showing overall distribution of predicted Z-scores (LN IC50) across tumor types. **f** Boxplots showing the distribution of GSVA scores of proliferation-related pathways across tumor types (Asterisks denote Wilcoxon rank sum test *p-value*). **g** Box plot depicting predicted Z-scores (LN IC50) of EGFR signaling pathway targeting drugs (Asterisks denote Wilcoxon rank sum test *p-value*). **h** Heatmap showing predicted Z-scores (LN IC50) for three ‘unseen’ drugs not present in GDSC, namely—BIC, APA, and ENZ. Color bars indicate tumor types and clusters as acquired through UMAP projections.

### Evaluation of models trained using TCGA patient tumor profiles

The Cancer Genome Atlas (TCGA) furnishes a large compendium of datasets spanning multiple cancer types with gene expression profiles of primary patient tumors and clinical response information. The clinical drug response data includes information about patient demographics and patient response for the drug administered. We aimed to perform classification by stratifying patients as responders who had a complete or partial response and non-responder for the patients who had clinically progressive or stable disease. To make the best use of the patient gene expression profiles and clinical drug response data, we used AutoML provided by open-source H2O.ai (https://docs.h2o.ai/h2o/latest-stable/h2o-docs/automl.html)36 in R to build a classifier. The summary of the dataset used is provided in supplementary data **(Supplementary Fig. 7)**. As discussed before, for CCLE bulk RNA-seq cell lines data, we integrated our processed TCGA pathway score matrix **(see methods)** with the vector embeddings of 139 unique drugs that were sourced from TCGA clinical drug response data. This matrix constituted 3108 patient-drug combinations in rows and 1427 features entailing 1327 pathway score vectors and vector embeddings of drug compounds of size 100 each, forming a set of explanatory variables to predict drug response as obtained from TCGA drug response data. For classification, we split this dataset and used 80% of the dataset for training the AutoML models, and 20% of the dataset was used for testing purposes. Our training dataset comprised of 2486 drug-patient combinations and the testing dataset consisted of 622 drug-patient combinations. AutoML is a platform that enables users to automatically train and tune multiple models by specifying a maximum number of models, thus automating machine learning workflows. We specified 20 maximum models to be trained with five fold cross validation. This resulted in 22 models, of which 20 models spanned across families of models, including GBM, XGBoost, DeepLearning, DRF, XRT models, and two stacked ensemble models. The two ‘stacked ensemble models’ were automatically trained and correspond to — one based on ‘all prior models’ that have been trained, while the other one is based on ‘each family’s best model’^36^. We evaluated the performance of the test dataset on these 22 trained models and found the ‘stacked ensemble model’, that was trained based on ‘all prior models’ outperform all others with an AUC=0.96, AUCPR= 0.98, and Gini=0.91 (**Fig. 4a**). Other evaluation metrics are reported in the supplementary data **(Supplementary Table S1)**. We used the best model for all subsequent prediction tasks. We also reported cross-validation performance metrics for these models **(Supplementary Table S2)**. Additionally, TCGA data have recorded survival information for patients. Using this information, we conducted survival analysis on the test dataset comprising 622 patients by stratifying patients based on the median value of the predicted probability of response. As shown in **Fig. 4b**, patients in group 1 who had a probability of response greater than the median value had better survival outcomes (*P* < 0.0001). We estimated the 5-years survival probability of group 1 as 0.774 and group 2 as 0.349. Then we sought to traverse the relevance of cancer stages in influencing the significant survival for group 1. For this, we visualized the frequency of cancer stages across different cancer types for group 1 and group 2. We observed that for the majority of cancer types, stage information is not known for most patients. Notably, in group 1, BRCA cancer type has the highest number of patients (n=34) belonging to stage iia, however, only 8 patients belonged to this stage in group 2. In our test dataset, we have a total of 42 BRCA individuals in stage iia, with 80.95 % of these patients falling into group 1 and 19.04 % falling into group 2 **(Supplementary Fig. 8)**. Our results are concordant with previous studies that early detection of cancer is associated with improved overall survival^37^. Further, **Supplementary Fig. 9** shows confusion matrices for group 1 and group 2 depicting the concordance of predicted and actual drug responses in patients. Next, we evaluated our model using a dataset for which pre and post-treatment RNA-seq profiles and clinical response information are available^38^. We predicted the drug response of RAF inhibitor (dabrafenib) and MEK inhibitor (trametinib) in three pre-treatment and matched post-lapse BRAF-mutant RAF/MEK inhibitor-resistant melanoma patients. **Fig. 4c** shows the journey of patient 1. The patient was diagnosed with Stage IIIC melanoma. After 1 year of initial treatment, the patient again showed signs of recurrent disease and underwent pre-treatment biopsy and intensity-modulated radiation therapy (IMRT). This patient exhibited both BRAF V600E and V600K mutations and was recruited into a Dabrafenib and Trametinib phase I/II study. After three months, the patient was removed from the study due to the development of resistance. Then the patient was treated with anti-PDL1 antibody for 4 months until progression and subsequently treated with four cycles of Ipilimumab. The patient died of his condition approximately nine months after the stoppage of RAF and MEK inhibitors. The journeys of other patients are documented in supplementary data (**Supplementary note 1)**. Wagle, Nikhil, et al. reported the presence of MEK2^Q60P^ mutation, BRAF Splice Isoform, and BRAF amplification in the post-treatment patient one, patient two, and patient three, respectively, as revealed through whole-exome sequencing (WES) and RNA-seq. These alterations seem to be a potential cause of conferring resistance to RAF/MEK inhibitors in these patients^38^. Notably, our prediction results revealed a similar trend, whereby our probability of response of dabrafenib and trametinib in pre-treatment patient one and patient two is higher than post-treatment (**Fig. 4d, e**).

**Fig. 4.**
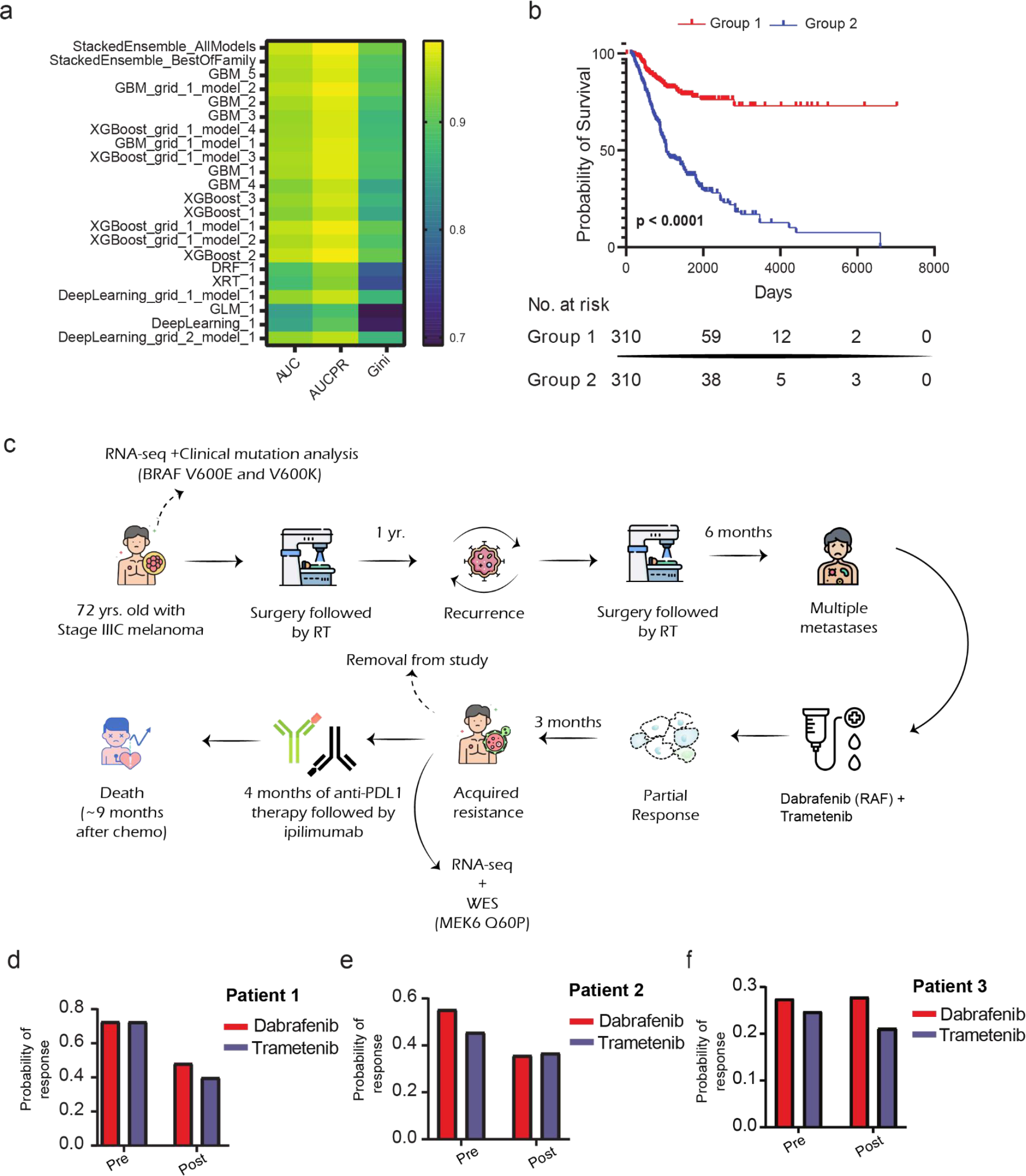
Evaluation of TCGA model efficiency. **a** Heatmap showing performance metrics, namely AUC, AUCPR, and Gini index for 22 AutoML trained models on the test dataset. **b** Survival analysis of a test dataset spanning multiple TCGA cancer types. Patients were classified into two groups based on the median value of the predicted probability of response with a P-value < 0.0001 (log-rank test). **c** Schematic representation of the journey of melanoma patient 1 from diagnosis of melanoma to initial treatments and enrolment into phase I/II study of dabrafenib and trametinib which resulted in initial regression of disease but ultimately the patient died after discontinuation of chemotherapy. **d-f** Bar plots showing the probability of predicted response for chemotherapy drugs dabrafenib and trametinib in patient 1, patient 2, and patient 3, respectively.

Furthermore, we were correctly able to predict these patients as responders. This aligns with the original annotations of the study as these patients are categorized as a partial response based on Response Evaluation Criteria In Solid Tumors (RECIST)^39^. In contrast, patient three was correctly predicted to be resistant to dabrafenib for both pre and post-treatment (**Fig. 4f**). This is concordant with the original study where this patient was categorized as a stable disease based on RECIST criteria.

## Discussion

Predicting drug response of cancer cells is of paramount importance for personalized oncology. In this study, we designed a deep neural network (DNN) model to predict the drug response by leveraging pathway activity scores derived from gene expression profiles of cell lines from CCLE and corresponding vector embeddings of drug compounds that were sourced from GDSC. Due to the explicit use of pathway enrichment scores, our model also highlights underpinning biological mechanisms that contribute to drug resistance. Furthermore, our pathway-based prediction approach allowed us to accurately infer cell fates upon treatment from single cell expression data. This can open doors for drug response prediction at sub-clonal resolution using tumour scRNA-seq data.

We evaluated the DNN model predictions by showing a reasonable correlation between the experimental Z-scores (LN IC50) for LNCaP cells used in the training data set and the model-predicted Z-scores. Further, we noted that *PTEN* negative LNCaP cell line was predicted to be sensitive to PI3K/mTOR signaling targeting drugs. Notably, previous studies have broadly suggested that increased PI3K/AKT/mTOR signaling is associated with sensitivity to PI3K inhibitors in LNCaP cell lines and other PTEN null cancer cell lines. Furthermore, mTOR inhibitors might be effective against PTEN null tumors^40^. With our predictive model, we could spot the sensitive phenotype of LNCaP cell lines to PI3K/mTOR inhibitors. This aligns with the previous findings in the LNCaP cell line.

With the confidence gained from cell line based validations, we further explored how drug sensitivity prediction changes upon drug treatments, drug-responsive and resistant states in LNCaP cells and xenografts. We sought to explore how these findings align with the known biology surrounding these treatments and states. In the presence of androgens, LNCaP cells were predicted to be more sensitive to cancer therapeutics that target highly proliferative cells. This is an expected result since androgens drive proliferation in AR-positive prostate cancer cell lines and xenografts^34,35^. AR antagonists are the primary treatment option for metastatic prostate cancer, which on a molecular level antagonize cellular androgen response pathways. With AR antagonist treatments, LNCaP cells were predicted to have pronounced similarities and differences for ENZ, APA, and BIC treatments, mirroring the complex biology underlying the treatment responses. The strongest reversal by an AR antagonist of DHT conferred sensitivity was observed for drugs targeting the PI3K/mTOR pathway, with ENZ showing the most profound effect. In the absence of androgens in all conditions, LNCaP was predicted to be resistant to drugs targeting DNA replication and genome integrity, namely irinotecan, cisplatin, and talazoparib. While ENZ treatment was uniquely predicted to increase the DHT conferred sensitivity to these drugs, BIC and APA partially decreased the DHT conferred sensitivity **(Supplementary Fig. 5b, c)**. These differential effects are important to be considered for combinatorial therapy in a given context of treatments and treatment resistance with other cancer therapeutics.

To further test the applicability of the drug sensitivity predictions, we have used data derived from an internally generated and biologically well-annotated prostate cancer xenograft model, following the progression from early androgen-responsive to CRPC, and then to ENZ treatment responsive and ENZ resistant states. The sensitivity predictions highlight changing vulnerabilities of the tumors in different stages of progression and treatment. Notably, we observed that ENZR tumors are predicted to develop new susceptibility to novel therapeutics providing a new window of opportunity for new therapeutic strategies. For instance, our model predicted sensitivity to specific drugs targeting the EGFR signaling pathway in case of ENZ treatment. While prostate tumors in an androgen replete setting are predicted to be highly resistant to EGFR targeting drugs, a distinct vulnerability is developed during progression to ENZR. As these drugs targeting the EGFR pathway are approved for cancer treatment in other cancer types, this may facilitate the clinical evaluation of combination therapy in prostate cancer patients treated with ENZ or those that have developed ENZR. Previous studies have suggested that a combination of ENZ and EGFR targeting inhibitors might be an effective therapeutic strategy in overcoming ENZ resistance^41^. Further laboratory and preclinical studies into the molecular mechanism behind our predicted sensitivity pattern are needed to confirm these findings.

Additionally, our AutoML model, trained on TCGA tumor RNA-seq data, revealed substantially improved overall survival in patients, predicted for partial or complete drug response. Further, testing of the TCGA model using the external BRAF-mutant melanoma dataset resulted in clinically relevant predictions for the patient samples. As expected, the probability of response for the patients classified in the partial response category was higher than post-treatment due to acquired resistance to the first line of therapy. This suggests that top drugs predicted by our method, for example, cyclophosphamide, cisplatin, might serve as alternative therapies in combination with other drugs to overcome acquired resistance.

There are some technical limitations of our work. In TCGA clinical drug response data, the same patients have been treated with multiple drugs, so we have treated them individually keeping the transcriptome identical.

Overall, this is among the first works connecting bioinformatic predictions of drug response to clinically explainable observations, both in *in vivo* and *in vitro* settings. Further, to our knowledge this is the first focused study on prostate cancer, investigating the potential of tumor bulk RNA-seq data in predicting drug response.

### Methodology

#### Prostate cancer cell lines and culture

The human prostate cancer cell lines LNCaP, DuCaP, VCaP, DU-145, and PC-3 were cultured in Phenol-red free RPMI medium supplemented with 5% fetal bovine serum (FBS) in a humidified incubator at 37°C and 5% CO2 and were harvested for RNA extraction during their exponential growth phase.

#### *In vitro* LNCaP cell line treatments

The human prostate cancer cell line LNCaP (#CRL-1740™ clone FGC, purchased from ATCC) was seeded into Phenol-red free RPMI medium supplemented with 5% fetal bovine serum (FBS) and cultured for 72 hr in a humidified incubator at 37°C and 5% CO2. This was followed by a 48 hr incubation in androgen-depleted conditions using medium + 5% charcoal-stripped serum (CSS). Treatment with the androgen targeting drugs (ATTs) enzalutamide (10uM), bicalutamide (10uM), and apalutamide (10uM) was performed for 48 hr, either in the absence or presence of 10 nM dihydrotestosterone (DHT, dissolved in EtOH).

#### Establishment of LNCaP xenografts

For the *in vivo* tumor progression study, xenografts were established by subcutaneous injection of one million LNCaP cells into the flank of male NOD-SCID mice. At a tumor size of 200 square mm, mice were either surgically castrated or received mock surgery for the PRE-CX group. Tumors from the PRE-CX group were harvested when the tumor size reached 1000 square mm. Tumors from the POST-CX (post castration) group were harvested 1-week post castration when serum PSA (Prostate-Specific Antigen) reached a nadir. Tumors from the CRPC (Castrate-Resistant Prostate Cancer) group were harvested once the tumor size reached 1000 square mm post castration. For the ’ENZ’ groups, daily treatment with 10 mg/kg ENZ commenced as serum PSA began to rise post castration. Tumors were harvested either at PSA nadir (ENZS) while on enzalutamide treatment or when the tumor size had reached 1000 square mm despite enzalutamide treatment (ENZR).

#### RNA extraction, library preparation, and bulk RNA-sequencing

For mRNAseq, total cellular RNA was extracted using the Norgen RNA Purification PLUS kit #48400 (Norgen Biotek Corp., Thorold, Canada) according to the manufacturer’s instructions, including DNase treatment. RNA quality and quantity were determined on an Agilent 2100 Bioanalyzer (Agilent Technologies, Santa Clara, USA) and Qubit®. 2.0 Fluorometer (Thermo Fisher Scientific Inc, Waltham, USA). Library preparation and sequencing was done using the ’Illumina TruSeq Stranded mRNA Sample Prep Kit (strand-specific, polyA enriched, Illumina, San Diego, USA) with an input of 500 ng - 1 ug total RNA (RIN>8), followed by paired-end sequencing with a read length of 100-150 bp and yielding about 30-60 M read pairs per sample.

RNAseq raw data was processed through a custom-designed pipeline. Raw reads were assessed with FastQC^42^, then trimmed using TrimGalore^43^, followed by alignments to the human genome (GRCh38 / hg38) and transcriptome (Ensembl.v.99 / Gencode.v.33, Jan-2020) using the STAR^44^ aligner and read quantification with RSEM^45^. For xenograft samples, STAR alignment was done against a chimeric human+mouse reference (mouse: GRCm38 / mm10, Gencode.v.M24 / Ensembl.v.99, Jan-2020), followed by RSEM read quantification. TPM values from the RSEM output were used for GSVA scoring.

#### Gene expression data of cancer cell lines

To predict and investigate anti-cancer drug response measured by Z-score of half-maximal inhibitory concentration (LN IC50), we used publicly available TPM (transcript per million) normalized RNA-seq gene expression profiles of 1019 Cancer Cell Line Encyclopedia (CCLE)^7^ cell lines quantified using the RSEM (RNA-Seq by Expectation-Maximization) software. The corresponding drug response information for the cell lines was sourced from the GDSC2 dataset of the Genomics of Drug Sensitivity in Cancer (GDSC) database^8^. In the GDSC2 dataset, some drug-cell line pairs have multiple LN IC50 measurements. In such cases, we averaged the LN IC50 values to avoid ambiguity. Then, LN IC50 values of all the drugs were transformed into Z-scores by using the formula (x-μ)/σ. The variable x represents LN IC50 values for a drug across cell lines, whereas μ & σ, the mean and standard deviation respectively. The RNA-seq gene expression profiles of 550 CCLE cell lines overlapping with the GDSC2 dataset cell lines were used for the model training. This matrix contained 57820 Ensembl Gene IDs which were converted into official gene symbols using gencode.v19.genes.v7_model.patched_contigs.gtf annotation file. This resulted in multiple Ensembl gene IDs corresponding to some of the individual gene symbols. We considered the average expression in such cases. At this stage, our expression matrix comprised 54301 genes and 550 cell lines. This matrix was subjected to log2 transformation with the addition of a pseudo count of 1.

#### Tumour RNA-seq data

Analogous to cell lines, on tumour mRNA sequencing data from TCGA, we modeled drug-response in terms of responder and non-responder. TCGA RNA-seq data was downloaded from the Broad GDAC firehose^46^ encompassing 33 tumor types. We used Illumina HiSeq RNA-seq v2 data processed at the gene level using RSEM^45^. The clinical drug response information for the patient samples was fetched from the NCI Genomic Data Commons portal^47^. Drug names and response information were collected from clinical metadata and rectified for typographical and spelling errors and to harmonize commercial names and molecular drug names. We categorized complete response and partial response patients as responders. Patients with clinically progressive and stable diseases were marked as non-responders. RNA-seq gene expression profiles of cancer types with clinical response data for fewer than two patients were excluded. At this stage, the filtered data contained gene expression profiles for 29 cancer types. Then for individual cancer types, scaled estimates from gene-level RSEM files were transformed into TPM by multiplying with a factor of 1e6 followed by log2 transformation with the addition of pseudo count 1. For the patients with identical TCGA barcodes, the ones with less non-zero gene expression were selected for downstream analysis.

#### Drug descriptor data

We obtained drug response information for 192 compounds from the GDSC2 dataset for the 550 cell lines in the CCLE dataset and clinical response information for 215 compounds for 1517 TCGA patients. The chemical structure information for these molecular compounds was retrieved in terms of a simplified molecular-input line-entry system (SMILES) using PubChemPy^24^. However, SMILES were not available for all the molecular compounds. As a result, we ended up with SMILES of 173 and 139 compounds for 550 CCLE cell lines and 1443 unique TCGA patients, respectively. The SMILES2Vec python tool was used to convert these SMILES into vector embeddings by utilizing data of embeddings trained on Pubchem and embedding of size 100^25^.

#### Pathway activity scores

To train models, we used pathway activity scores. For computing pathway activity scores, we used the Gene Set Variation Analysis (GSVA)^15^ R software package to log2(TPM + 1) gene expression matrix and gene set file from Molecular Signatures Database (MSigDB)^48^ with min.sz set as 5. We used the c2 collection of canonical pathways (MSigDB.CP.v.6.1) consisting of 1329 gene sets. We integrated the pathway score matrix with the vector embeddings of the drug features. Our final CCLE cell line training dataset constituted 80056 drug/cell line combinations in rows and 1429 features entailing 1329 pathways and drug features of vector size 100 for each molecular compound as the explanatory variable and Z-scores as the response variable in columns. For TCGA patient data, gene expression profiles of individual cancer types were transformed into pathway scores. The GSVA scores of the samples where drug response information was available in each cancer category were merged based on common pathways. The final matrix consisted of 3108 drug/patient combinations and 1427 features (pathways & drug descriptors), and the response variable (responder=1, non-responder=0).

#### Training models using CCLE RNA-seq cell line dataset

For drug response prediction in gene and pathway space, formulated as a regression task, we used state-of-the-art machine learning methods. We split the CCLE training dataset into an 80% training set and a 20% test set and trained ten bootstrapped regression models for each method using 70% samples as training data. The Random forests were constructed using the ranger R package^49^ with 500 trees. ElasticNet was implemented using the glmnet R package^50^. A deep neural network (DNN) was trained using the Keras platform. DNN was modeled with one input layer of as many features detected in the dataset, followed by two hidden layers of sizes 512 and 64, respectively, using RELU as an activation function. Dropout layers were added to avoid the overfitting of models with a dropout rate of 0.2. The ADAM optimizer and Mean squared error as loss function was used. Models were trained using 500 epochs and a batch size of 50. For the model genes as features, the DNN model was trained with an input layer of 600 features and with one hidden layer of 30 using RELU as an activation function. Like for pathway space models, ADAM optimizer and Mean squared error as loss function was used in gene space as well. Final predictions were computed as an average of all the predictions from the bootstrapped models.

#### Training models using TCGA RNA-seq patient profiles

To predict drug response in terms of responder and non-responder, formulated as a classification problem, we utilized the H2O AutoML^36^ framework in R for training the TCGA dataset. We split this dataset in the ratio of 80% for training and 20% for testing. This 80% training dataset was subjected to the h2o.automl() function with five-fold cross-validation and a maximum number of models of 20. This resulted in automated training of multiple machine learning models, including Deep learning, DRF, GBM, GLM, XGboost, XRT, and stacked ensemble models.

#### Survival analysis on TCGA test dataset

The processed TCGA dataset consisted of 3108 drug-patient combinations. We conducted survival analysis on a 20% TCGA test dataset composed of 622 drug-patient entities from 517 unique patients. We stratified the samples using the median value of the predicted probability of the response and computed survival along with 5 year survival probability.

#### Performance Metrics

We used two accuracy metrics to measure the performance of our models for the regression task: the coefficient of determination (R^2^) and Pearson correlation (ρ). R^2^ was calculated using the R caret^51^ package. For the TCGA dataset, we used MSE, RMSE, LogLoss, Mean Per-Class Error, AUC, AUCPR, and Gini index.

#### Imputation of missing features

We used an impute^52^ package from R to impute if any pathway features are missing in the input test dataset using the nearest neighbor-based averaging approach.

#### Validating CCLE cell line trained models using scRNA-seq data of cell lines

The scRNA-seq cell line pre-QC UMI count dataset comprising 30314 genes and 56982 cells encompassing 207 cell lines was obtained from Kinker, G. S. et al study^30^. This expression matrix was processed for quality control using an R script (data_processing.R) from the same study. At this stage, we were left with 53299 cells spanning 198 cell lines. This matrix was transformed into TPM by scaling with a factor 1e6 and dividing the UMI count of genes by the total UMI count of the sample. The UMI counts are independent of gene length bias^53^. We log2-transformed this normalized matrix with the addition of a pseudo count of 1. The gene expression values of the same cell lines were averaged. This matrix was converted into pathway scores using GSVA with default parameter settings. Our final validation set consisted of 17279 drug cell line combinations entailing 116 cell lines that overlapped with the cell lines featured in the GDSC2 dataset and tested against 173 drugs.

#### Predicting the response of paclitaxel drug in MDA-MB-231 single cells using CCLE cell line trained models

Lee et al. dataset was obtained by processing FASTQ files of MDA-MB-231 cells from SRA id SRP040309. We align FASTQ files using STAR aligner^44^ with GRCh37 reference genome and GRCh37 GTF file from Ensembl (release 75)^54^. To estimate gene expression, we used HTSeq-count^55^. The HTSeq generated read counts were transformed into TPM values by dividing raw counts by gene length, then scaling with a factor of 1e6 and dividing by total read counts in the sample. We discarded the genes for which gene length was not retrieved by the EDASeq R package^56^. This was followed by the conversion of Ensembl gene IDs to official gene symbols using Homo_sapiens.GRCh37.75.gtf annotation file. We retained genes having a TPM of at least 1 in at least 10% of samples. This matrix was subjected to log2 transformation with the addition of pseudo count 1. We averaged gene expression values of five samples each of naive, stressed, and cells more sensitive to paclitaxel respectively and converted them into pathway scores using GSVA. We predicted drug response for treatment-naive and population of cells more sensitive to paclitaxel in terms of Z-score of LN IC50 values.

#### Validating TCGA data trained models using BRAF-mutant melanoma patient profiles

We used publicly available Reads Per Kilobase of transcript per million mapped reads (RPKM) RNA-seq profiles of BRAF mutant melanoma patients for drug response prediction. This dataset comprises six paired patients for which RNA-seq was performed before treatment and after treatment with RAF or RAF+MEK inhibitors during disease progression. This dataset includes therapy response information as well. This dataset is referred to as Wagle, Nikhil, et al. dataset^38^. For our analysis, we transformed RPKM normalized RNA-seq profiles to TPM by dividing each RPKM value by the sum of the RPKM values for all genes in the sample and multiplying by one million^57^. The TPM normalized matrix was log2 transformed with the addition of a pseudo count of 1. The resulting matrix was transformed into pathway scores using GSVA. For the three patients for whom information on the mechanism of acquired resistance to dabrafenib/trametinib is available, we predicted a therapeutic response to dabrafenib and trametinib.

#### Code availability https://github.com/SmritiChawla/PrecOnco

### Author contributions

DS conceived the study with CCN. CCN and BGH contributed to the experimental design of prostate cancer study. SC conducted data analyses with assistance from SP, PR and NB. AJ, AA, AG, ND trained machine learning models. DS, AM and Jayadeva supervised the development of machine learning models. AR supervised and performed significant parts of data analysis and data interpretation. ML contributed to the RNA-seq pipeline of prostate cancer datasets, data interpretation and analysis. ER contributed in experimental design and animal work for LNCaP xenografts study, RNA extraction and sample QC. GA assisted in biological interpretation of results. DS, CCN, BGH supervised the entire study. All authors substantially contributed for manuscript writing and reviewing.

---

**Supplementary Fig. 1.**
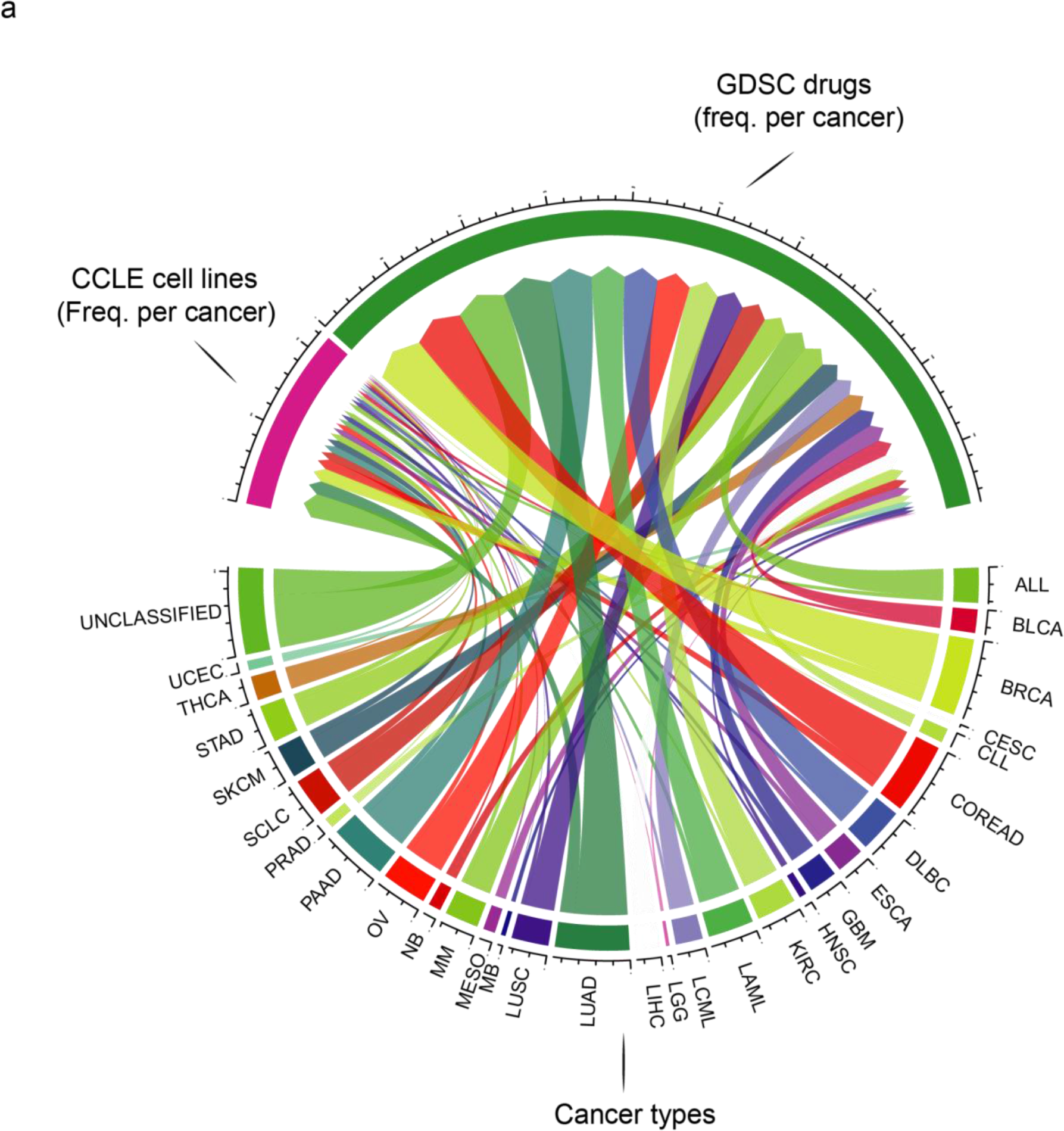
Overall summary of the CCLE and GDSC dataset used for training the Deep Neural Network (DNN). **a** Chord diagram showing the frequency of cell lines (n=550) from the CCLE database along with the frequency of tested drugs (n=173) from the GDSC database spanning 29 classified cancer types used for the training dataset.

**Supplementary Fig. 2.**
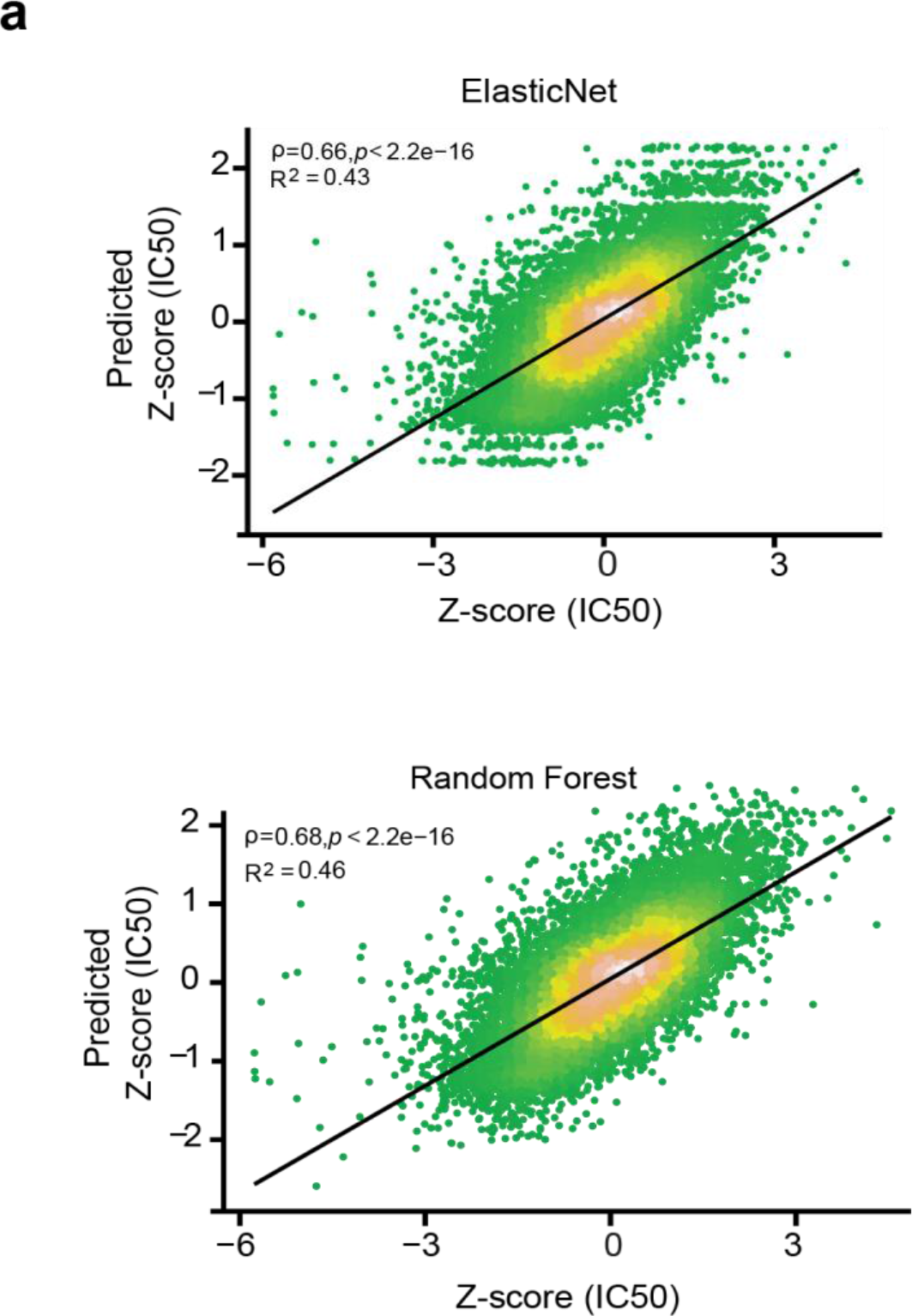
Performance evaluation of different machine learning models trained using CCLE bulk RNA-seq cell lines in pathway space. **a** Scatter plot of observed and predicted Z-score (IC50) showing prediction performance of Elastic net and Random Forest. We calculated Pearson correlation (ρ) and coefficient of determination (R^2^) for evaluation.

**Supplementary Fig. 3.**
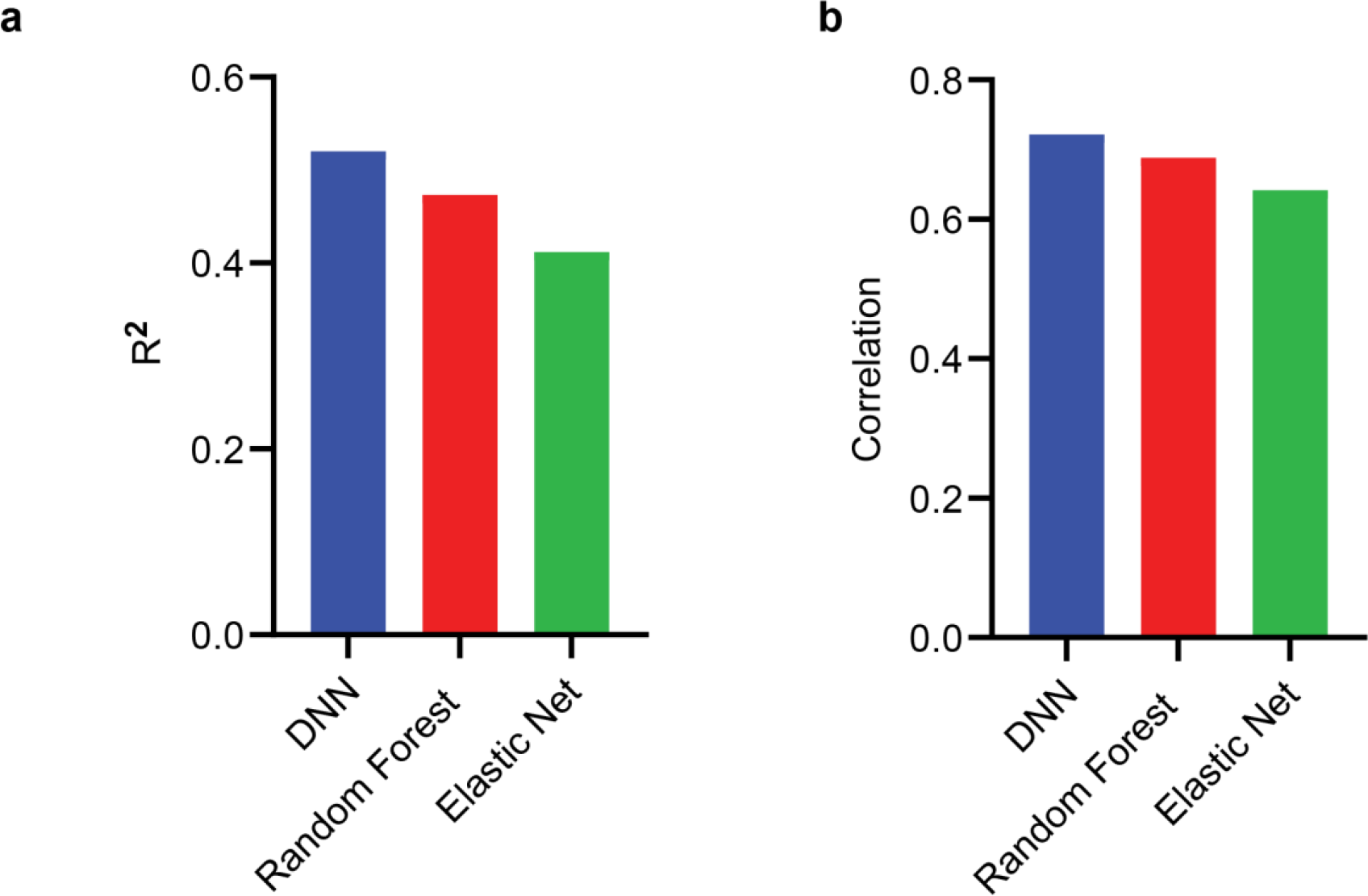
Performance evaluation of different machine learning models trained using CCLE bulk RNA-seq cell lines gene space. **a** Barplots showing coefficient of determination (R^2^). **b** Barplots showing Pearson correlation (ρ).

**Supplementary Fig. 4.**
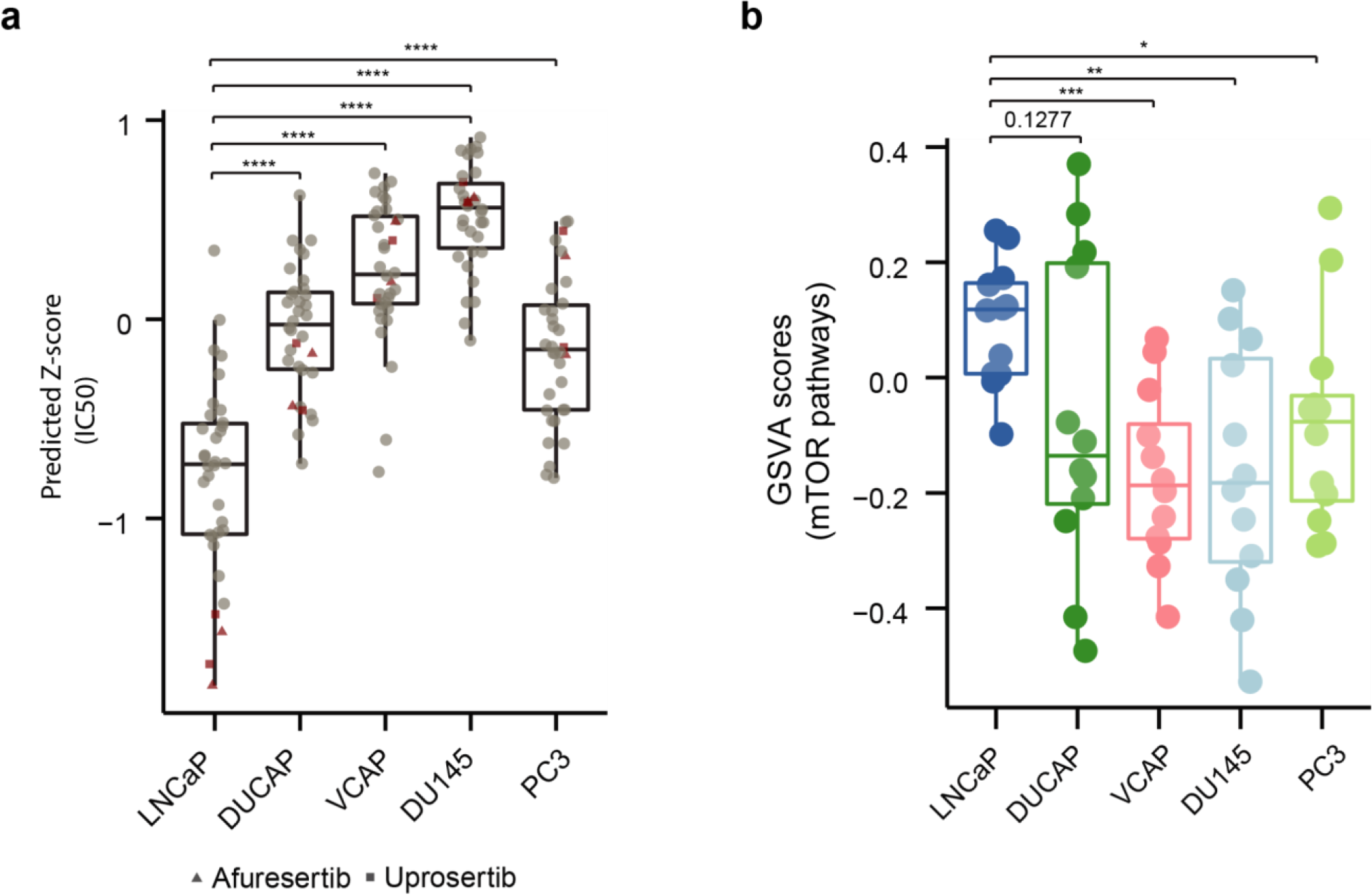
LNCaP cell line sensitivity to drugs targeting mTOR signaling. **a** Box plots depicting the distribution of drug response prediction of mTOR/PI3K signaling targeting drugs. LNCaP cell line, in particular, was predicted to be more sensitive to these drugs, with afuresertib and uprosertib having the most profound effect (Asterisks denote Wilcoxon rank sum test *p-value*). **b** Box plots showing the distribution of GSVA scores of mTOR-related pathways highlighting increased expression of these pathways in the LNCaP cell line (Asterisks denote Wilcoxon rank sum test *p-value*).

**Supplementary Fig. 5.**
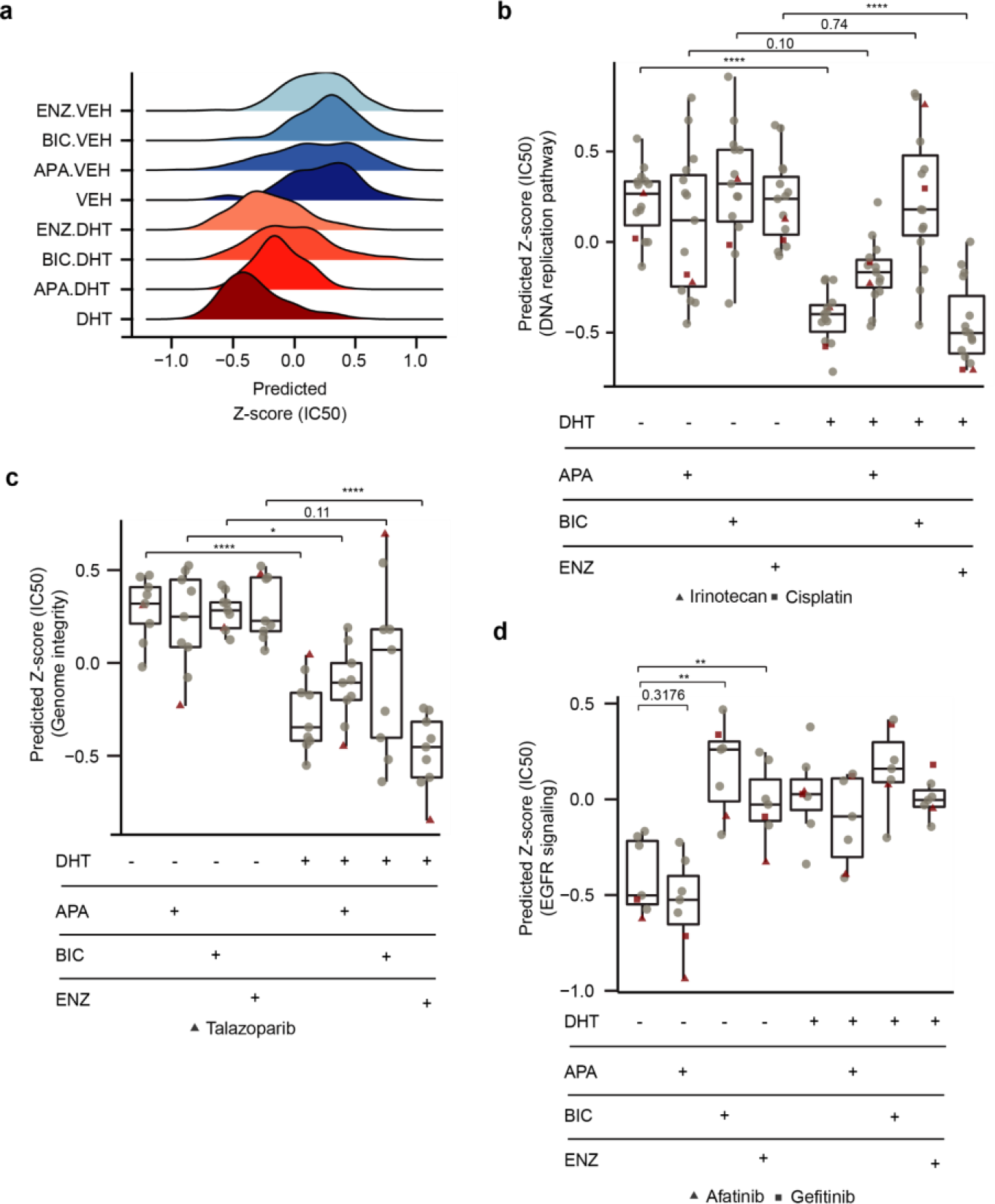
Drug response prediction and analysis in LNCAP cells under different treatment conditions. **a** Ridge plot showing the overall distribution of predicted Z-score (IC50) of 155 drugs tested against prostate cancer cell lines in the GDSC2 dataset across the treatment conditions. **b** Boxplot depicting predicted Z-score (IC50) of drugs targeting DNA replication pathway explicitly highlighting irinotecan and cisplatin (Asterisks denote Wilcoxon rank sum test *p-value*). **c** Boxplot showing predicted Z-score (IC50) of drugs targeting genome integrity pathway specifically pinpointing talazoparib (Asterisks denote Wilcoxon rank sum test *p-value*). **d** Boxplot of predicted Z-score (IC50) of the drugs targeting EGFR signaling across the treatment conditions specifically highlighting afatinib and gefitinib drugs (Asterisks denote Wilcoxon rank sum test *p-value*).

**Supplementary Fig. 6.**
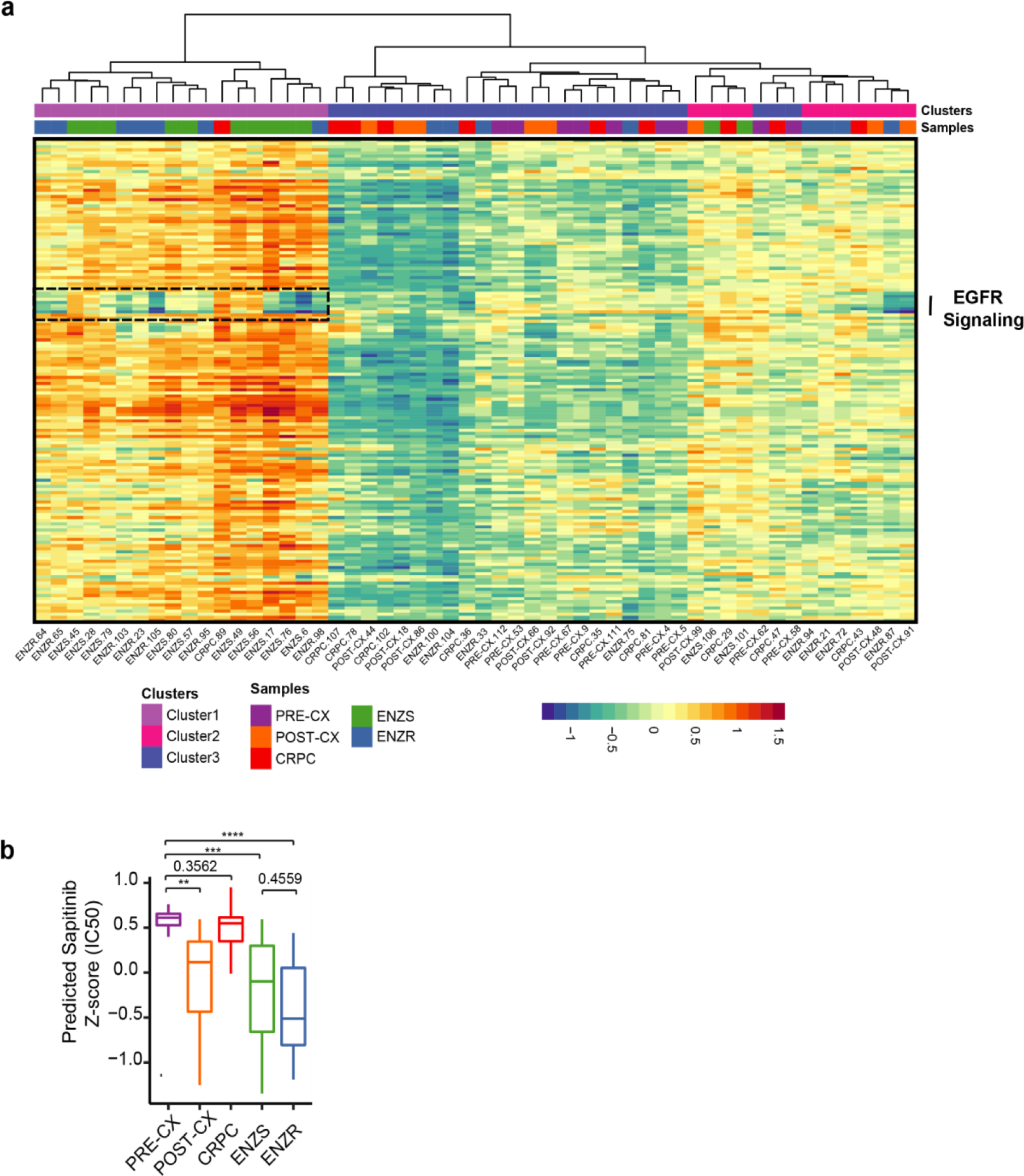
Predictions in LNCAP xenografts under different treatment conditions. **a** Visualization of predicted Z-scores (IC50) in 54 xenograft tumor samples for 155 GDSC drugs. Heatmap showing the sensitivity of some ENZ treated tumors to ‘EGFR signaling’ targeting drugs. **b** Boxplot of predicted Z-scores (IC50) of sapitinib across the tumor types revealing the highest sensitivity of this drug for ENZR tumors (Asterisks denote Wilcoxon rank sum test *p-value*).

**Supplementary Fig. 7.**
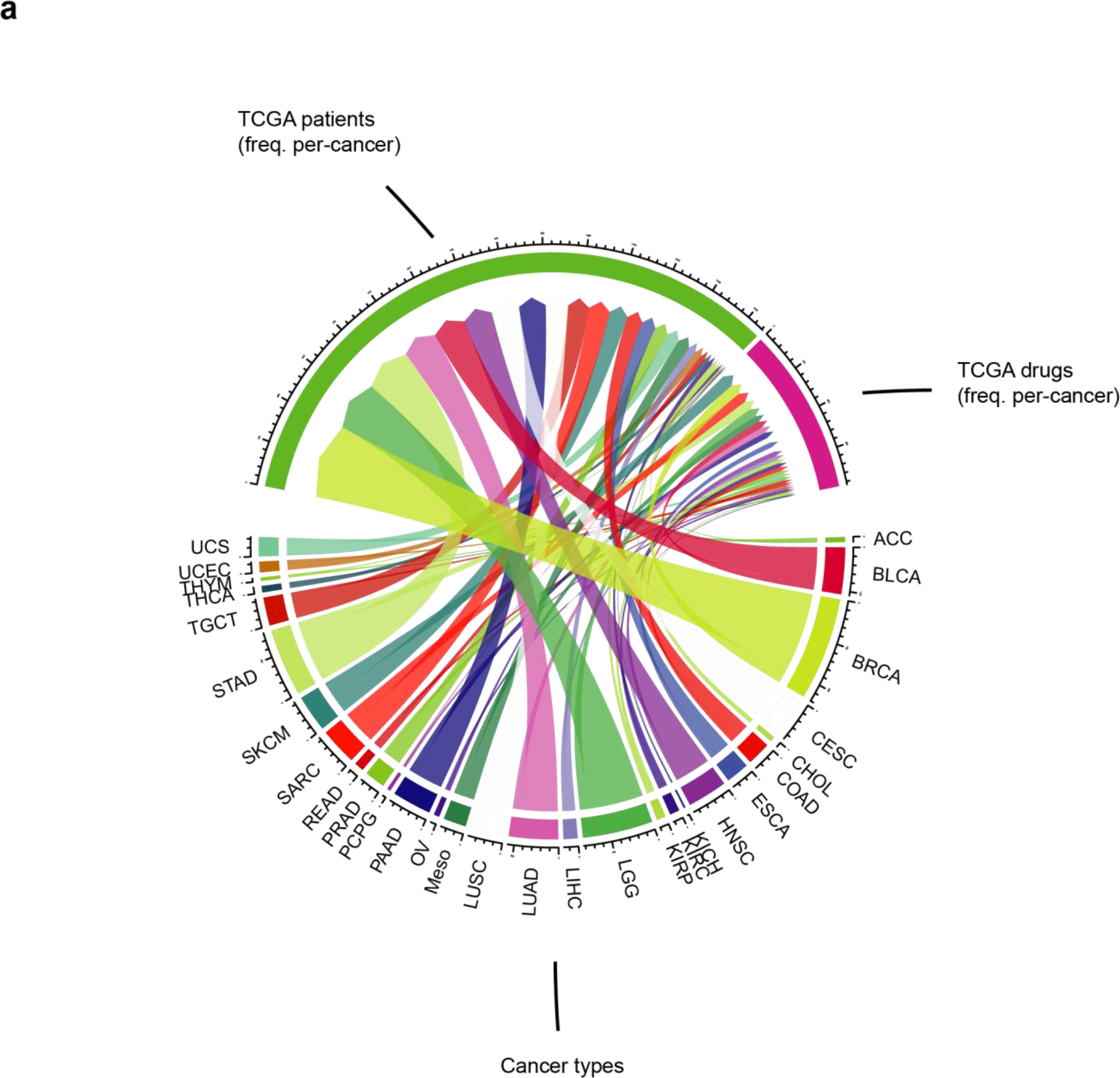
Description of the dataset used for training AutoML models. **a** Chord diagram showing the frequency of patients (n=1443) from the GDAC firehose database along with the frequency of tested drugs (n=139) from TCGA clinical response data spanning 29 classified cancer types used for the training dataset.

**Supplementary Fig. 8.**
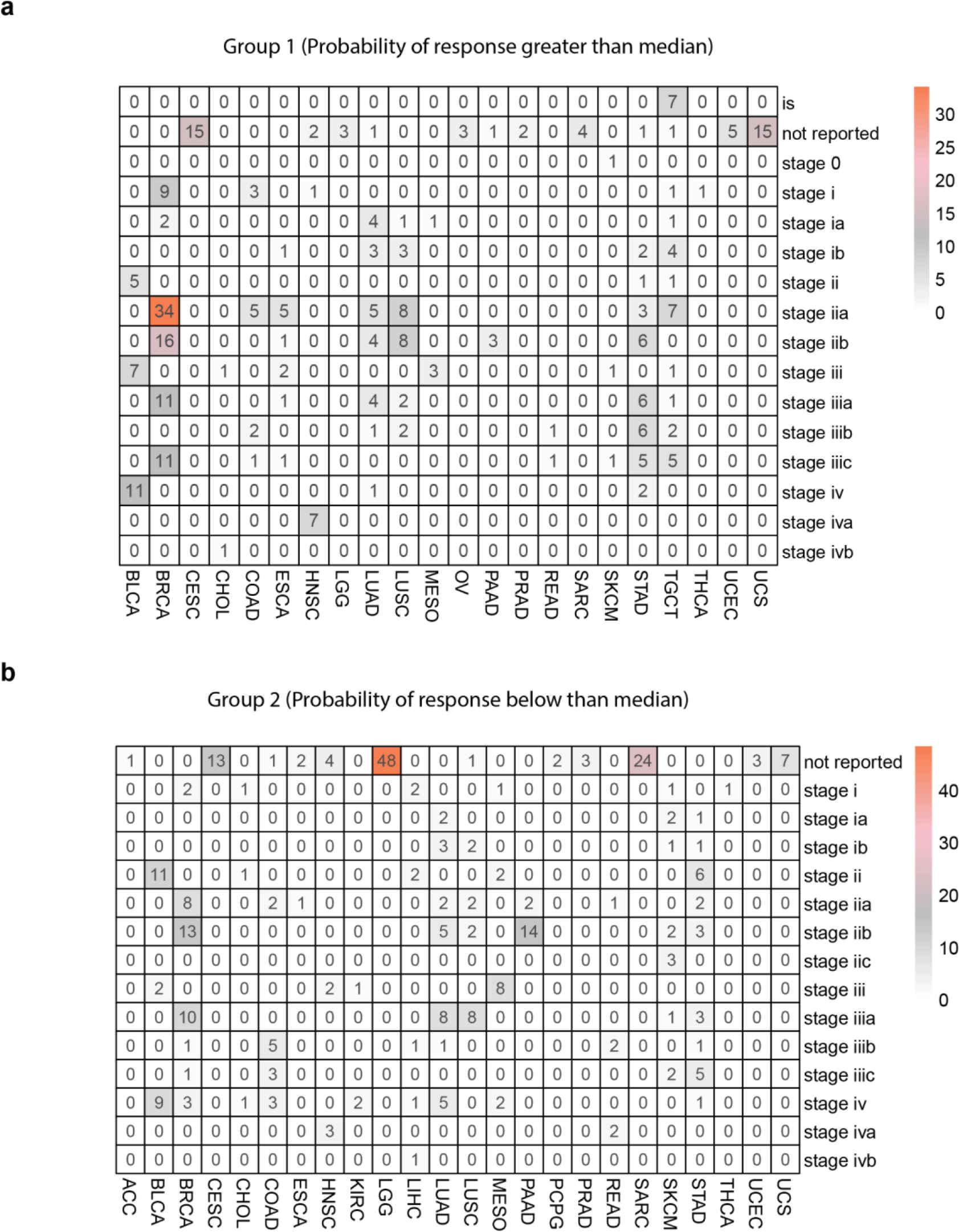
Visualization of frequency of stages across different cancer types for TCGA test dataset. **a & b** Heatmaps showing the frequency distribution of stages across cancer for group 1 and group 2, respectively, stratified based on the median value of the probability of response as predicted through classifier trained on TCGA dataset. We denoted the frequency of stage with 0 if stage is unknown.

**Supplementary Fig. 9.**
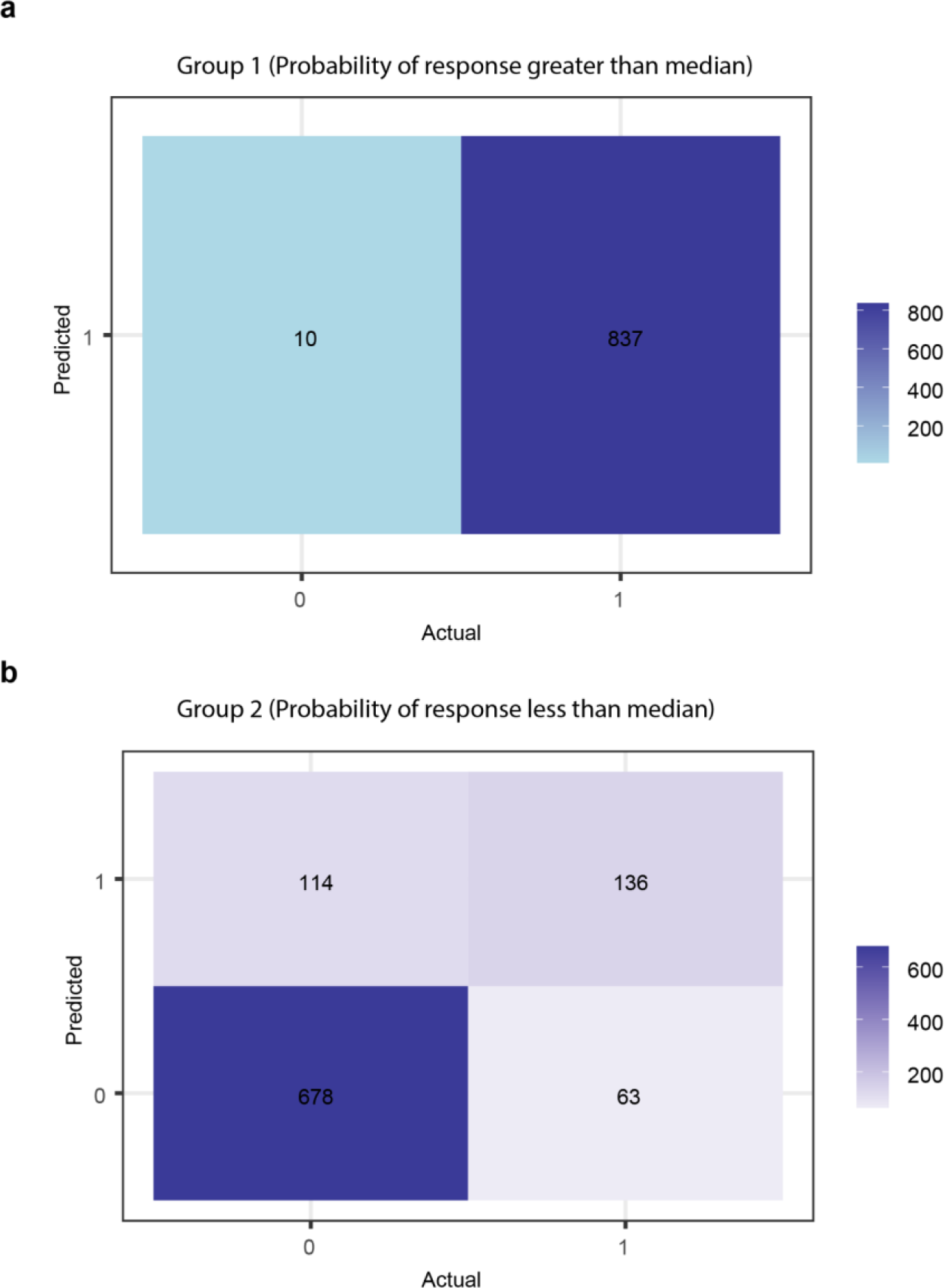
Visualization of confusion matrices for classification results of TCGA test dataset. **a** Heatmap for confusion matrix showing classification results for group 1. We stratified patients into group 1 and group 2 based on the median value of the probability of response as predicted through a classifier trained on TCGA dataset. Our median value of 0.80 yielded only patients belonging to class 1 (responder) in group 1. **b** Heatmap for confusion matrix showing classification results for group 2. The actual class depicts treatment response information from TCGA clinical metadata files. The treatment response information is categorized as responder (label=1) for complete response and partial response of patients. While clinical progressive disease and stable disease in patients represent non-responder categories (label=0).

**Supplementary Table 1.** Performance summary of AutoML models on the bulk RNA-seq TCGA patient test dataset.

| **Models** | **MSE** | **RMSE** | **Log Loss** | **Mean per class error** | **AUC** | **AUCPR** | **Gini** |
| --- | --- | --- | --- | --- | --- | --- | --- |
| StackedEnsemble_AllModels_AutoML_20210812_215305 | 0.08 | 0.28 | 0.27 | 0.1 | 0.96 | 0.98 | 0.91 |
| StackedEnsemble_BestOfFamily_AutoML_20210812_215305 | 0.09 | 0.29 | 0.29 | 0.1 | 0.95 | 0.97 | 0.9 |
| GBM_5_AutoML_20210812_215305 | 0.09 | 0.3 | 0.31 | 0.11 | 0.94 | 0.96 | 0.89 |
| GBM_grid_1_AutoML_20210812_215305_model_2 | 0.09 | 0.3 | 0.29 | 0.1 | 0.95 | 0.97 | 0.9 |
| GBM_2_AutoML_20210812_215305 | 0.1 | 0.31 | 0.31 | 0.11 | 0.94 | 0.96 | 0.89 |
| GBM_3_AutoML_20210812_215305 | 0.1 | 0.32 | 0.33 | 0.15 | 0.94 | 0.96 | 0.87 |
| XGBoost_grid_1_AutoML_20210812_215305_model_4 | 0.1 | 0.31 | 0.31 | 0.12 | 0.94 | 0.96 | 0.88 |
| GBM_grid_1_AutoML_20210812_215305_model_1 | 0.1 | 0.32 | 0.33 | 0.13 | 0.94 | 0.97 | 0.88 |
| XGBoost_grid_1_AutoML_20210812_215305_model_3 | 0.09 | 0.3 | 0.3 | 0.11 | 0.94 | 0.97 | 0.89 |
| GBM_1_AutoML_20210812_215305 | 0.1 | 0.31 | 0.32 | 0.13 | 0.94 | 0.97 | 0.89 |
| GBM_4_AutoML_20210812_215305 | 0.11 | 0.33 | 0.34 | 0.14 | 0.93 | 0.96 | 0.85 |
| XGBoost_3_AutoML_20210812_215305 | 0.1 | 0.32 | 0.32 | 0.15 | 0.93 | 0.96 | 0.87 |
| XGBoost_1_AutoML_20210812_215305 | 0.11 | 0.33 | 0.34 | 0.17 | 0.93 | 0.95 | 0.85 |
| XGBoost_grid_1_AutoML_20210812_215305_model_1 | 0.08 | 0.29 | 0.28 | 0.13 | 0.95 | 0.97 | 0.91 |
| XGBoost_grid_1_AutoML_20210812_215305_model_2 | 0.09 | 0.3 | 0.3 | 0.11 | 0.95 | 0.96 | 0.89 |
| XGBoost_2_AutoML_20210812_215305 | 0.08 | 0.29 | 0.28 | 0.1 | 0.95 | 0.97 | 0.91 |
| DRF_1_AutoML_20210812_215305 | 0.14 | 0.38 | 0.44 | 0.22 | 0.89 | 0.93 | 0.78 |
| XRT_1_AutoML_20210812_215305 | 0.15 | 0.38 | 0.45 | 0.23 | 0.88 | 0.93 | 0.76 |
| DeepLearning_grid_1_AutoML_20210812_215305_model_1 | 0.11 | 0.33 | 0.52 | 0.12 | 0.93 | 0.96 | 0.87 |
| GLM_1_AutoML_20210812_215305 | 0.15 | 0.39 | 0.47 | 0.24 | 0.85 | 0.89 | 0.69 |
| DeepLearning_1_AutoML_20210812_215305 | 0.16 | 0.41 | 0.53 | 0.23 | 0.85 | 0.9 | 0.7 |
| DeepLearning_grid_2_AutoML_20210812_215305_model_1 | 0.1 | 0.31 | 0.42 | 0.12 | 0.93 | 0.95 | 0.87 |

**Supplementary Table 2.**
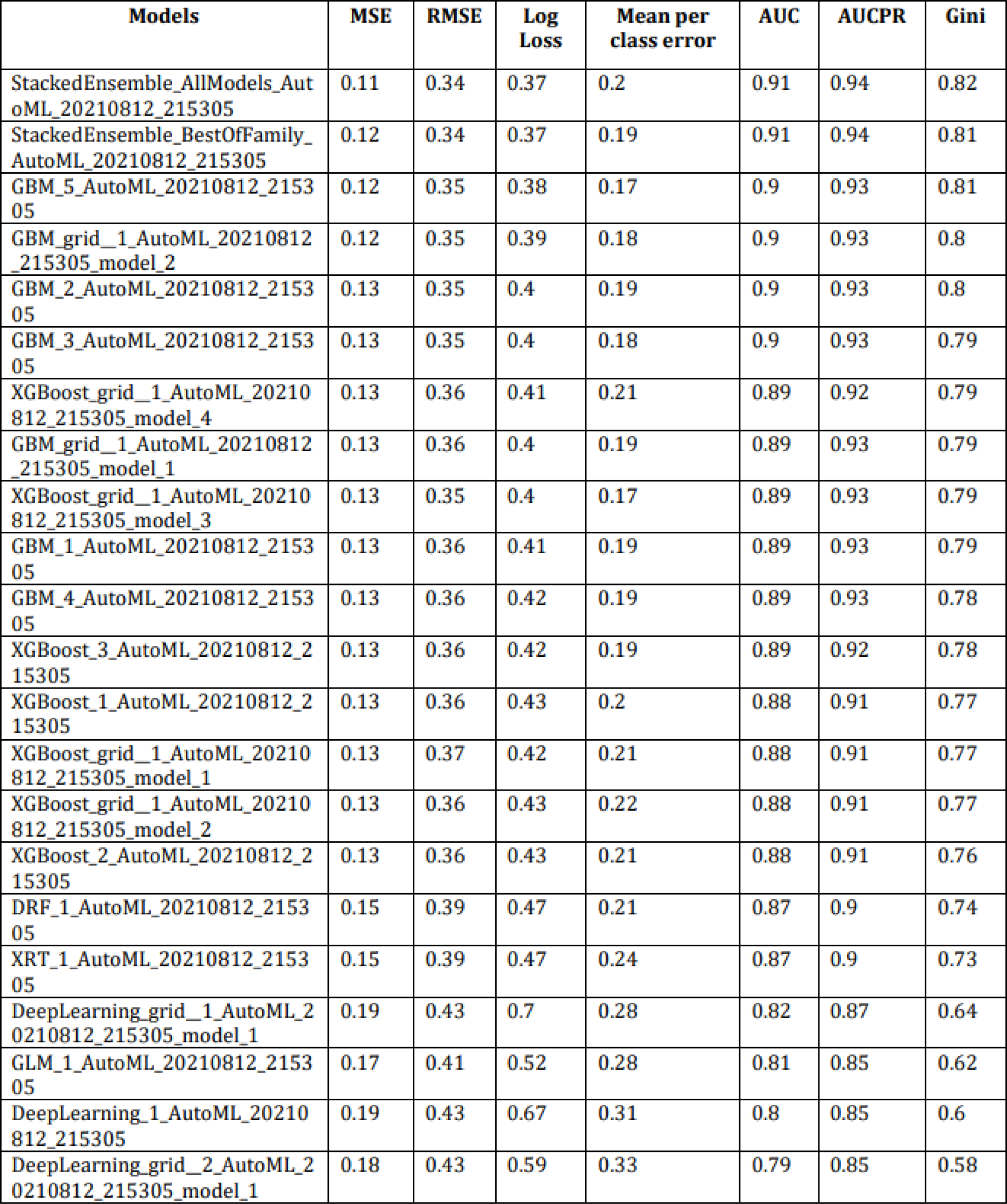
Performance summary of AutoML models on bulk RNA-seq TCGA patient profiles training dataset (cross-validation)

## Supplementary data

Supplementary note 1: Clinical characteristics of melanoma patients Patient 2

Patient 2, a 48-year old man initially diagnosed with stage IB melanoma developed extensively metastatic melanoma after five years of initial diagnosis, confirmed by a pleural biopsy (pre-treatment). Further, clinical mutation analysis revealed the presence of BRAF V600E mutation. The patient was subjected to first-line treatment of Dabrafenib and Trametinib which showed partial response but after 3 months, routine scans revealed significant disease progression. The potential cause of resistance to therapy was the presence of BRAF splice variant as detected by RNA-seq and whole exome sequencing (WES) in post-treatment tumors but not in pre-treatment tumors. The patient died six months after being diagnosed with metastatic disease^38^.

Patient 3

Patient 3, a 42-year old man underwent surgery and lymph node dissection of stage IIIC melanoma of the left thigh. The patient had a BRAF V600E mutation as revealed by clinical mutational analysis. After six months of surgery, the patient was subjected to first-line therapy of dabrafenib and trametinib. But nearly after one year the patient developed progressive disease and the potential underlying cause of acquired resistance to therapy was the presence of BRAF amplification in post-treatment tumors as depicted by WES. The patient was given Ipilimumab for a short time but died after four cycles and three months after discontinuation of chemotherapy^38^.

